# GPR4 Deficiency Alleviates Intestinal Inflammation in a Mouse Model of Inflammatory Bowel Disease

**DOI:** 10.1101/059014

**Authors:** Edward J. Sanderlin, Nancy R. Leffler, Kvin Lertpiriyapong, Qi Cai, Heng Hong, Vasudevan Bakthavatchalu, James G. Fox, Joani Zary Oswald, Calvin R. Justus, Elizabeth A. Krewson, Elizabeth A. O’Rourke, Li V. Yang

## Abstract

GPR4 is a proton-sensing G protein-coupled receptor that can be activated by extracellular acidosis. It has recently been demonstrated that activation of GPR4 by acidosis increases the expression of numerous inflammatory and stress response genes in vascular endothelial cells (ECs) and also augments EC-leukocyte adhesion. Inhibition of GPR4 by siRNA or small molecule inhibitors reduces endothelial cell inflammation. As acidotic tissue microenvironments exist in many types of inflammatory disorders, including inflammatory bowel disease (IBD), we examined the role of GPR4 in IBD using a dextran sulfate sodium (DSS)-induced colitis mouse model. We observed that GPR4 mRNA expression was increased in mouse and human IBD tissues when compared to control intestinal tissues. To determine the function of GPR4 in IBD, wild-type and GPR4-deficient mice were treated with 3% DSS for 7 days to induce acute colitis. Our results showed that the severity of colitis was decreased in GPR4-deficient DSS-treated mice in comparison to wild-type DSS-treated mice. Clinical parameters, macroscopic disease indicators, and histopathological features were less severe in the DSS-treated GPR4-deficient mice than the DSS-treated wild-type mice. Inflammatory gene expression, leukocyte infiltration, and isolated lymphoid follicle (ILF) formation were reduced in intestinal tissues of DSS-treated GPR4-null mice. Collectively, our results suggest GPR4 provides a pro-inflammatory role in IBD as the absence of GPR4 ameliorates intestinal inflammation in the acute DSS-induced IBD mouse model.

## Introduction

The pH-sensing G protein-coupled receptors (GPCRs) have emerged as a new class of receptors that are involved in sensing both local and systemic pH changes. Subsequently, these receptors have been implicated in various disease states associated with dysregulated pH homeostasis such as cancer, ischemia, metabolic acidosis, and inflammation ^1–3^. Family members of the pH-sensing GPCRs include GPR4, TDAG8 (GPR65), OGR1 (GPR68), and G2A (GPR132) ^1, 2, 4–9^. These receptors are capable of sensing protons in the extracellular milieu by the protonation of several histidine residues on their extracellular domains ^4, 8, 10^. GPR65, GPR68, and GPR132 are predominately, though not exclusively, expressed on leukocytes and provide various roles in the exacerbation or amelioration of a diverse set of diseases associated with inflammation and acidosis ^1, 11, 12^. GPR4, reciprocally, is highly expressed in vascular endothelial cells (ECs) and blood vessel rich tissues such as the lung, kidney, heart, and liver ^9, 13–16^. Recently, GPR4 has been shown to mediate EC inflammatory responses to acidosis and is central for leukocyte-endothelium interaction ^17, 18^.

In response to extracellular acidosis (increased extracellular proton concentration), GPR4 has been reported as a pro-inflammatory mediator in a variety of ECs ^17, 18^. Both isocapnic and hypercapnic acidosis have been demonstrated to activate GPR4 and induce an inflammatory response in three types of primary endothelial cells, including human umbilical vein endothelial cells (HUVECs), human pulmonary artery endothelial cells (HPAECs), and human lung microvascular endothelial cells (HMVEC-Ls) ^17, 18^. The GPR4 mediated inflammatory response to acidosis encompasses the induction of adhesion molecules such as E-selectin (SELE), vascular cell adhesion molecule 1 (VCAM-1), and intercellular adhesion molecule 1 (ICAM-1) in ECs and subsequently increases the functional adhesion of leukocytes *in vitro* ^17, 18^. In addition to adhesion molecules, GPR4 activation in ECs increases the expression of chemokines such as CCL20, CXCL2, and IL-8 (CXCL8) involved in the recruitment and activation of leukocytes ^17, 18^. Furthermore, GPR4 activity stimulates the induction of COX-2, NF-κB pathway genes, and stress responsive genes in ECs under acidic conditions. These results collectively describe GPR4 as pro-inflammatory through increasing leukocyte-EC adhesion and subsequent extravasation into inflamed tissues ^17, 18^. Therefore, GPR4 could potentially provide a role in the inflammatory response for host defense and the removal of pathogens or apoptotic cells in various tissues by the recruitment of leukocytes. If inflammation is not properly resolved, however, GPR4 could exacerbate inflammatory disorders.

Recently, a family of imidazo pyridine derivatives has been identified as exhibiting antiinflammatory functions in ECs by reducing pro-inflammatory cytokine secretion, adhesion molecule expression, and leukocyte-EC adhesion through the inhibition of GPR4 ^17, 19, 20^. In addition to chemical antagonists of GPR4, similar results were observed with use of siRNA inhibitors specifically targeting GPR4 expression ^18^. Moreover, it has been shown that the expression of the GPR4 gene can be stimulated in ECs by inflammatory stresses such as cytokines (TNF-α) and reactive oxygen species (H_2_O_2_) ^15^, which commonly exist in inflammatory bowel disease.

Inflammatory bowel disease is characterized by chronic, aberrant mucosal inflammation of the gastrointestinal tract ^21^. There are two distinct disease subsets in which IBD can take form, namely, Crohn’s disease (CrD) and ulcerative colitis (UC). The exact etiology of IBD is unknown, but a complex interaction between immunologic, environmental, microbiome, and genetic constituents is believed to contribute to the disease onset and continued progression.
Both CrD and UC have distinct, yet overlapping clinical and histopathological features that are a result of altered mucosal homeostasis. The production of cellular metabolic byproducts contributes to an acidic inflammatory mucosal loci in IBD ^22^. Indeed, an acidic inflammatory locus is a hallmark of chronically inflamed tissue as numerous studies have shown that local tissue pH below 7.0, and sometimes even below 6.0, is detected in inflammatory diseases and alters cellular functions ^22–27^. In addition to tissue acidosis in the gut, reports indicate that the lumen of the colon is more acidic in patients with IBD than patients without IBD ^28, 29^. As a result, host vasculature, leukocyte infiltrates, and stromal cells often function within an acidic tissue microenvironment and can in turn modulate the inflammatory response ^22^.

Inflammation in IBD is a conglomerate of gut associated pathologies, but one particular pathological hallmark is a hyper-dysregulated vasculature inflammatory response in the gut ^30^. Host vasculature is critical in mediating the extent of inflammation and subsequent tissue damage resulting from chronic inflammation. The inflammatory response requires the active passage of leukocytes such as neutrophils, monocytes, and lymphocytes to the site of inflammation through host vasculature to the gut mucosa. EC adhesion molecules and chemokines facilitate leukocyte complementary binding for firm adhesion and subsequent extravasation from the blood vessel wall into tissue. The endothelium therefore functions as a gate; either barring or allowing the passage of inflammatory cells into inflamed tissue. Modulating the passage of leukocytes into tissue is an ideal target for IBD therapy. Currently, anti-adhesion biologics such as natalizumab and vedolizumab are used in the clinic for IBD patients ^31, 32^. Even though anti-adhesion therapies have proven efficacious in the clinical remission of IBD, there have been some limitations reported. For example, cases of progressive multifocal leukoencephalopathy (PML) have been observed in patients treated with natalizumab ^33, 34^.

We hypothesize that endothelial GPR4 expression functions as a “gatekeeper” in regulating the extent of leukocyte infiltration into the inflamed colon. In this study, we observed that GPR4 mRNA expression was increased in the inflamed colon of human IBD samples as well as in a DSS-induced IBD mouse model. Using GFP knock-in as a surrogate marker for GPR4 expression in GPR4 knockout (KO) tissues, we were able to observe GFP expression in endothelial cells of intestinal microvessels, arteries, veins, high endothelial venules (HEVs), and histiocytes. GPR4 expression, however, was not detected in lymphatic vessels. GPR4 deficiency reduced the overall inflammation parameters used to gauge the extent of IBD severity. Furthermore, inflammatory gene expression positively correlated to increased GPR4 mRNA expression in wild-type (WT)-DSS colon tissue, but was overall reduced in GPR4 KO-DSS tissues when compared to WT-DSS tissues. ILF development was also reduced in the inflamed colon of GPR4-deficient mice compared to WT mice with intestinal inflammation most likely due to the proposed role of GPR4 in regulating leukocyte extravasation through blood vessels. Altogether, our study has identified GPR4 as a potential regulator of intestinal inflammation and suggests that molecular responses to the acidic microenvironment in inflamed intestinal tissues may be a novel mechanism involved in IBD pathogenesis. A similar mechanism may also exist in other inflammatory disorders.

## Methods

### Dextran sulfate sodium (DSS)-induced acute IBD mouse model

All experiments were carried out in 9 week old male and female wild-type and GPR4-deficient mice. The mice were backcrossed to C57BL/6 background for 11 generations. The mice were maintained specific pathogen-free of exogenous murine viruses, ectoparasites, endoparasites, and *Helicobacter.* Mice were housed in an Association for Assessment and Accreditation of Laboratory Animal Care (AAALAC)-accredited facility under environmental conditions of a 12:12 light/dark cycle, temperature maintenance at 22 ± 1°C and relative humidity range of 30-70%. Mice were group housed in microisolator caging on corncob bedding and provided autoclaved tap water and pelleted diet (ProLab 2000, Purina Mills, St. Louis, MO) *ad libitum.* Colitis was induced by the addition of 3% (w/v) Dextran Sulfate Sodium Salt (DSS) [36,000-50,000 M.Wt, MP Biomedical, Solon, OH] to autoclaved drinking water. Mice were treated with 3% DSS or water for seven consecutive days, with a replenishment of 3% DSS or water every two days. Mouse body weight and clinical phenotypic scores were assessed daily during the treatment period and tissue was collected at the end of the treatment period. Animal studies were performed according to the randomized block experimental designs that can increase the power and reproducibility ^35^. All animal experiments were approved by the Institutional Animal Care & Use Committee of East Carolina University, Greenville, North Carolina and were in accordance with the *Guide for the Care and Use of Laboratory Animals* administered by the Office of Laboratory Animal Welfare, NIH.

### Clinical scoring

Colitis severity was quantified using the clinical phenotype parameters of weight loss and fecal score which were determined daily for each mouse. The fecal score was determined using the parameters of stool consistency and fecal blood (0= normal, firm, dry; 1 = formed soft pellet, negative hemoccult test; 2 = formed soft pellet with positive hemoccult test; 3 = formed soft pellet with visual blood; 4 = liquid feces with visual blood; 5 = no feces, only bloody mucus or empty colon upon necropsy). Presence of fecal blood was determined by the use of the Hemoccult Single Slides screening test (Beckman Coulter, Brea, CA).

### Collection of tissue for histology and molecular analysis

After the seven day treatment of DSS, mice were euthanized and the entire gastrointestinal tract was removed. The colon length was measured from anus to ileocecal junction, then detached from the cecum. The colon was then washed with phosphate buffered saline (PBS) to remove fecal matter. Six, five millimeter sections of the colon were resected commencing from the anus and moving toward the cecum and promptly snap frozen in liquid nitrogen for storage at a-80°C freezer for RNA analysis. The remaining colon tissue was fixed in 10% buffered formalin for further histological analysis. The cecum was also cleaned of all fecal matter and fixed in 10% buffered formalin for histological studies. The mesenteric lymph nodes were isolated. The lymph nodes most proximal to the cecum were used for size measurement and histological analysis. Lymph node length (L) and width (W) were measured to calculate the volume of each lymph node using the formula (L×W^2^) π/6. Once the measurements were taken, the lymph nodes were fixed using 10% buffered formalin and processed for histological analysis.

### Histopathological scoring

Two independent board certified pathologists, with expertise in animal or human pathology, analyzed colon and cecum 5μm sections stained with hematoxylin and eosin (H&E) in a blind fashion. Pathologists analyzed and scored sections for histopathological features commonly observed in IBD tissues. Scoring criteria and methodology were conducted as previously reported with minor adaptations ^36, 37^. Each pathologist used complementary, yet distinct scoring systems for histopathological analysis. Briefly, the criteria used by the veterinary pathologist (V.B.) for histopathological changes included a scoring system from one to four, wherein a score of four was most severe. The veterinary pathologist assessed and scored the degree of inflammation, epithelial defects, crypt atrophy, epithelial hyperplasia, and dysplasia. Each cecum and colon section was assessed for each parameter and the sum of each parameter was presented as total histopathological score per mouse ^36^. The scoring criteria used by the human pathologists (Q.C. and H.H.) included the individual assessment of each parameter including inflammation, area of leukocyte infiltration, crypt damage, and edema. The score for each parameter was multiplied by a factor corresponding to the degree of overall intestinal tissue involvement. The sum of all parameters for each mouse provided the total histopathological ^37^.

### Isolated lymphoid follicle quantification

Cecum and colon tissue was collected as described above and serial histological sections were stained with hematoxylin and eosin (H&E). Using a light microscope, cecum tissue sections were scanned using 4× and 10× objective lenses and isolated lymphoid follicles (ILFs) were counted. Colon sections were scanned from proximal to distal on longitudinal sections using 4× and 10× objective and ILFs were counted. The entire colon section was then measured in centimeters. ILF number is presented as ILFs per centimeter of colon section length. ILFs in the cecum are presented as ILFs per cecum in tissue sections.

### Immunohistochemistry (IHC)

Immunohistochemistry was performed on serial sections of 5-μm paraffin-embedded cecum, lymph node, and colon tissue sections. All colon, cecum, and mesenteric lymph node sections were de-paraffinized and hydrated from 100% ethanol to water followed by antigen retrieval using Tris-EDTA pH 9.0 with 0.1% Tween 20. Slides were incubated in antigen retrieval buffer for 18 minutes at 99 °C followed by blocking of endogenous peroxidase activity. Tissue sections used to analyze GFP expression in GPR4 KO mice were incubated with primary goat polyclonal against green florescent protein (GFP) (Abcam, ab6673, 1:1000, Cambridge, MA). The IHC system (Anti-goat HRP-DAB Cell and Tissue Staining Kit, R&D Systems, Minneapolis, MN) was used which employs a peroxidase-conjugated streptavidin as a colorogenic component. For IHC analysis of adhesion molecules E-selectin (CD62E) and VCAM-1, SuperPicture 3rd Gen IHC Detection system (Invitrogen, Waltham, MA) was employed. To block endogenous mouse IgG in blood serum, Mouse on Mouse blocking reagent (Vector Laboratories, Burlingame, CA) was used and followed by blockade with 10% normal serum. Primary antibodies against E-selectin/CD62E (Abcam, ab18981, 1:500, Cambridge, MA) or VCAM-1 (Abcam, ab134047, 1:1000, Cambridge, MA) were incubated overnight at 4°C and then recombinant secondary antibody incubation occurred followed by DAB (3,3′-diaminobenzidine) incubation. Slides were then dehydrated and mounted. Pictures were taken with a Zeiss Axio Imager A1 microscope.

### Real-time RT-PCR

Total RNA was isolated using IBI Scientific DNA/RNA/Protein Extraction Kit (MidSci) and 500-1000ng of RNA were reverse transcribed using SuperScript II reverse transcriptase (Invitrogen, Waltham, MA). TaqMan pre-designed primers-probe sets specific for target gene (Applied Biosystems, Foster City, CA) were used and are listed in supplementary information (SI Table 1). Real-time PCR was performed in duplicate with a program of 50°C for 2 min, 95°C for 10 min followed by 40 cycles of 95°C for 15 sec and 60°C for 1 min, and the data was acquired and analyzed using the ABI 7300 or ABI 7900HT real-time PCR thermocycler machines. Crohn’s and colitis cDNA arrays were purchased from Origene Technologies (catalog #CCRT102, Rockville, MD) and subjected to real-time PCR using specific primer-probes for human GPR4 and β-actin. The primer and probe used for human GPR4 has been previously described ^18^. The cDNA array contained 47 samples including 7 normal, 14 Crohn’s, and 26 ulcerative colitis intestinal samples from patients diagnosed with IBD.

### Statistical analysis

GraphPad Prism software was used to perform the statistical analyses. Differences between two groups (wild-type mice versus GPR4 knockout mice) were compared using the Mann-Whitney test or unpaired *t* test. Correlation of gene expression was determined by the linear regression analysis. *P* < 0.05 is considered as statistically significant.

## Results

### GPR4 deficiency attenuates the severity of colitis in the DSS-induced IBD mouse model

In order to determine the functional role of GPR4 in intestinal inflammation, we chemically induced intestinal inflammation in wild type (WT) and GPR4 KO mice. By day 7, WT mice treated with DSS (WT-DSS) lost nearly 16% of bodyweight on average (Fig. 1A). In comparison, GPR4 KO mice treated with DSS (GPR4 KO-DSS) had only a 7% reduction in bodyweight. The WT-DSS mice also had a fecal score more severe than GPR4 KO-DSS mice intermittently throughout the experiment (Fig. 1B). On day 7 of the experiment, mice were euthanized and the colon length was evaluated as an indicator of the degree of colonic inflammation/fibrosis induced by DSS. GPR4 KO-DSS mice had less colon shortening compared to WT-DSS mice (Fig. 1C). Additionally, mesenteric lymph nodes were isolated and measured to calculate the volume as an assessment of the response to inflammatory stimulation. GPR4 KO-DSS mice had a significant reduction in mesenteric lymph node expansion compared to WT-DSS mice (Fig. 1D). Collectively, the clinical phenotype of gut inflammation was less severe in GPR4 KO-DSS mice compared to WT-DSS mice indicating GPR4 is pro-inflammatory in IBD. The DSS-induced colitis disease model causes very severe acute intestinal inflammation and tissue damage ^38, 39^; subsequently, the partial recovery phenotype observed in the GPR4 KO-DSS mice is comparable to other studies showing an alleviation of the DSS-induced phenotype with mutant mice ^40–43^.

In conjunction with the clinical aspects of colonic inflammatory extent, severity was also assessed through histopathological analysis by both veterinary and human pathologists. Common features of IBD were evaluated and scored such as the degree of inflammation, area of leukocyte infiltration, edema, epithelium damage, hyperplasia, dysplasia, and crypt damage for both the cecum and colon. Both independent pathologists, using distinct methodologies, arrived at the same observation that GPR4 KO-DSS mice were less severe when compared to WT-DSS mice in both the colon and cecum (Fig. 2; SI:1). Of particular interest, the degree of inflammation and area of leukocyte infiltration were reduced spanning from cecum and colon tissues in the GPR4 KO-DSS mice compared to WT-DSS mice (Fig. 2C-D). Interestingly, the degree of reduction in histopathological features of GPR4 KO-DSS mice were significantly greater in the cecum when compared to colon (Fig. 2A-B). This observation could be due to the increased acidity in the cecum compared to the colon ^28, 44, 45^, thereby increasing GPR4 activity in the cecum.

**Figure 1.**
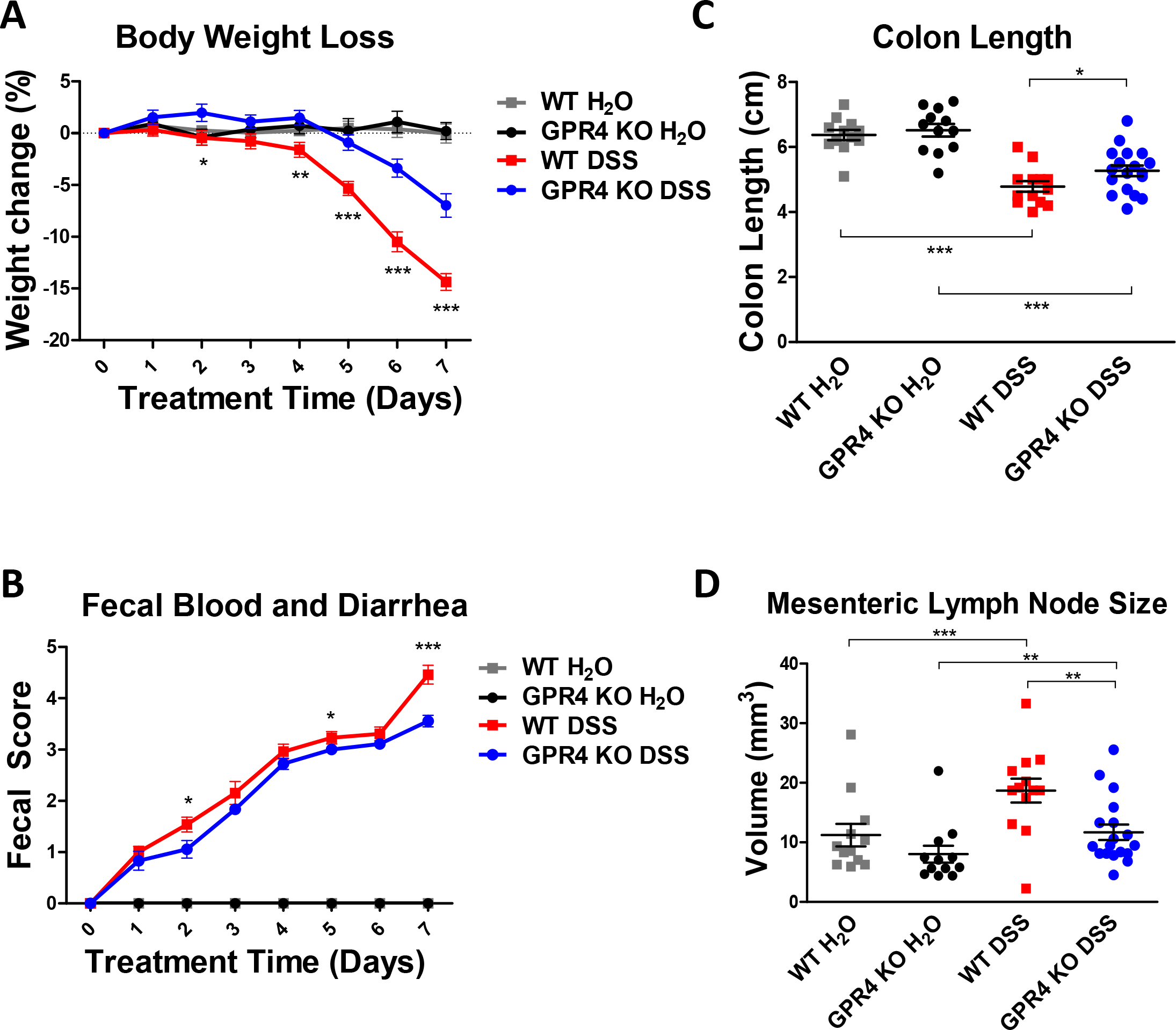
| Clinical phenotypes and macroscopic indicators. To assess the extent of DSS-induced IBD in mice, we measured several parameters to gauge the disease severity in mice. We observed GPR4 KO-DSS mice had reduced IBD severity when compared to WT-DSS mice. Clinical parameters of IBD severity, including (A) Body weight loss, (B) colon length, (C) fecal score and (D) mesenteric lymph node volume, were assessed in WT-control (n=12), WT-DSS (n=13), GPR4 KO-control (n=12), and GPR4 KO-DSS (n=18) mice. Each dot represents the data from an individual mouse. Data are presented as mean ± SEM and statistical analysis is between WT-DSS mice and GPR4 KO-DSS mice or as indicated in graph. (\**P* < 0.05, \*\**P* < 0.01, *** *P*< 0.001).

**Figure 2.**
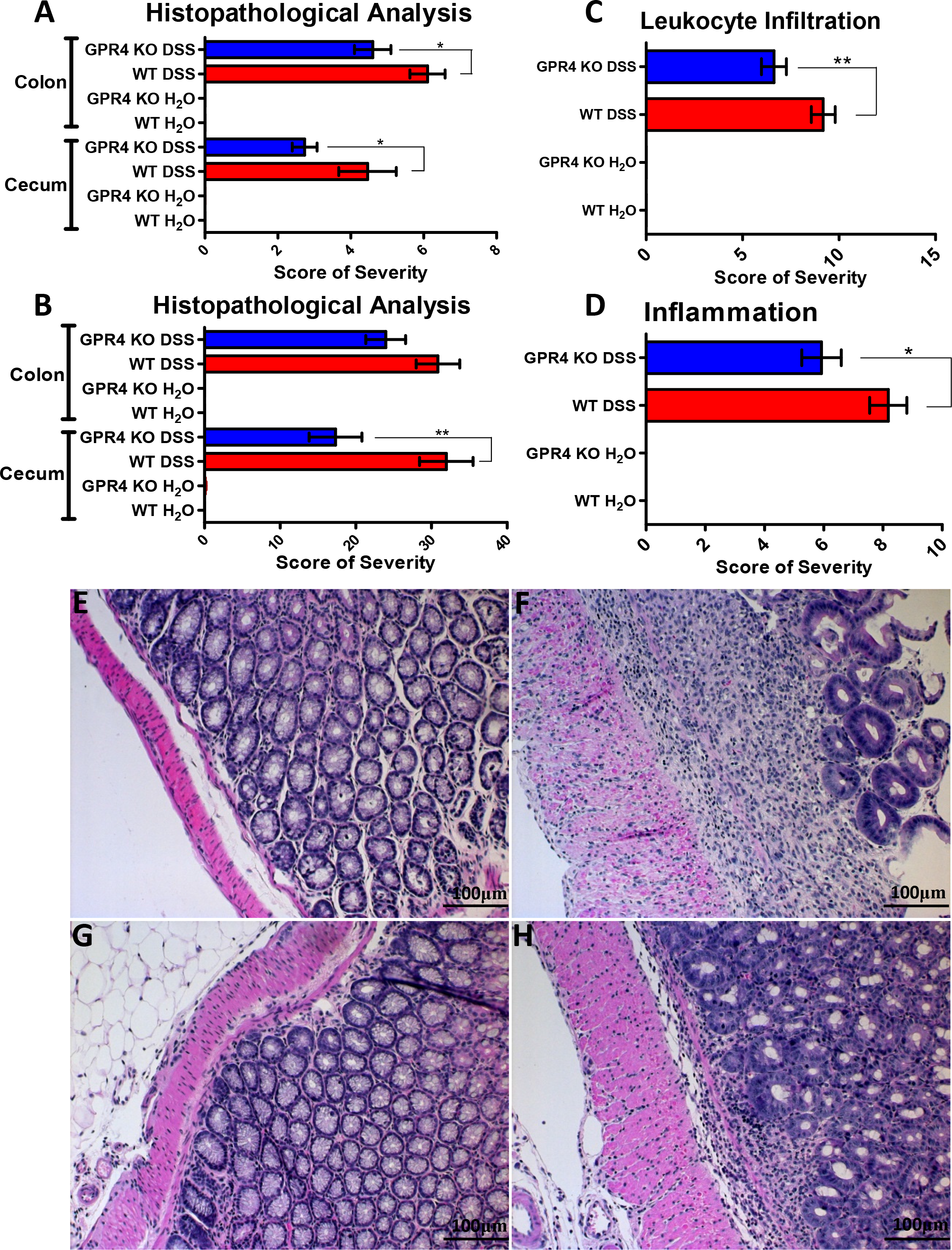
| Histopathological analysis of mouse colon and cecum. Histological features of IBD were examined to further assess the degree of disease activity in the mice by veterinary and human pathologists using complementary, yet distinct scoring systems. Overall, GPR4 KO DSS mice had reduced histopathological scores in cecum and colon when compared to WT DSS mice. Veterinary pathologist (A) and human pathologist (B, C, D) assessment of colon and cecum. Reduced leukocyte infiltration was observed spanning from cecum to colon in GPR4 KO DSS mice compared to WT DSS mice (C). Overall inflammation was reduced in GPR4 KO DSS mice compared to WT DSS mice in tissues spanning from cecum to colon (D). Representative H&E staining pictures of colon in WT control (E), WT DSS (F), GPR4 KO control (G), and GPR4 KO DSS (H). 20× microscope objective. Data are presented as mean ± SEM (\**P* < 0.05, \*\**P* < 0.01)

### Formation of lymphoid follicles in the colon and cecum is reduced in DSS-treated GPR4-null mice

Recently there has been growing interest in isolated lymphoid follicles (ILFs) and local gut immunity. ILFs predominately develop in the colon and are similar in structure and function as Peyer’s patches (PP) in the small intestine with the major difference between PP and ILFs being the inducible nature of ILFs in response to inflammatory stimuli ^46, 47^. Crosstalk between stromal cells, lymphoid tissue inducer cells (LTi), and immune cells (dendritic cells, T cells, B cells) are critical for ILF development and effector functions in the gut ^48^. Increased development of ILFs in the colon is associated with increased intestinal inflammation and tissue damage ^49^. In keeping with these reports, we observed a significant increase in ILF development in WT-DSS mouse colons when compared to WT control colons (4.0 ILFs/cm vs. 0.8 ILFs/cm of colon section, *P* < 0.001) (Fig. 3A). Similar results were observed upon evaluation of the cecum sections for ILFs (Fig. 3B; SI:2). The GPR4 KO-DSS mice, however, had no significant increase of ILF formation in both the colon and the cecum (Fig. 3; SI:2). The GPR4 KO-DSS mice had on average 1.1 ILFs per centimeter of colon section. ILFs could be observed spanning the proximal and distal sections of the colon. ILF density increased in areas of greater inflammation. As such, fewer ILFs were observed in the proximal regions of the colon compared to the distal colon where increased inflammation was visible. The GPR4 KO-DSS mice had an ILF number very similar to the WT-control mice in both colon and cecum. These results suggest GPR4 is critical for ILF formation in response to gut inflammation.

**Figure 3.**
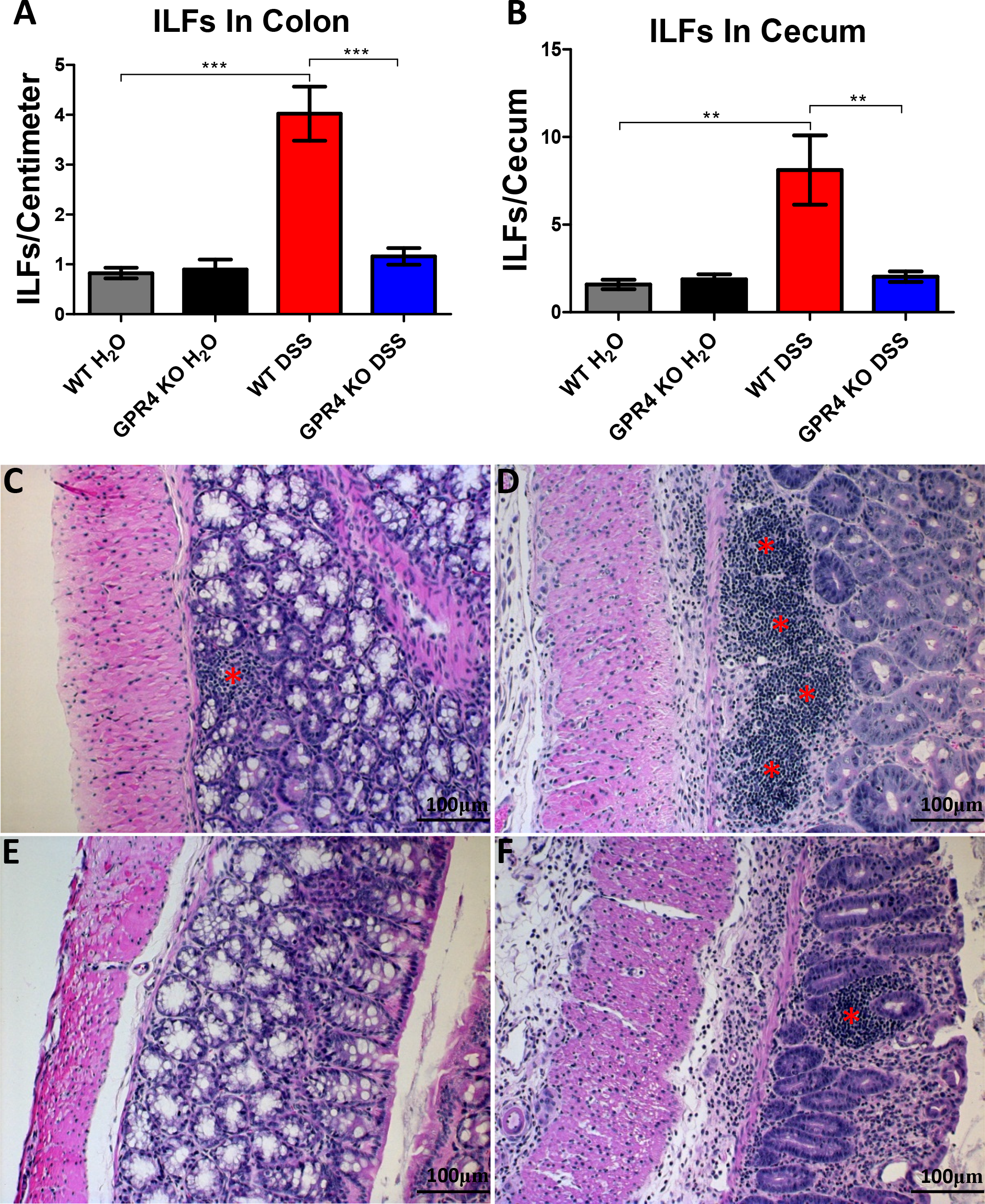
| Isolated lymphoid follicle (ILF) quantification and H&E staining of ILFs. GPR4 KO DSS mice had reduced ILF number in colon and cecum when compared to WT DSS mice. ILF quantification in WT and GPR4 KO colon (A) and cecum tissues (B). Representative H&E staining of ILFs in WT control (C), WT DSS (D), GPR4 KO control (E), and GPR4 KO DSS (F) colons. Red asterisks indicate representative ILFs in colon tissue. Data are presented as mean ± SEM (\*\**P* < 0.01, *** *P*< 0.001)

### GPR4 mRNA expression is increased in human and mouse IBD intestinal tissues compared to normal intestinal tissues

To further characterize the involvement of GPR4 in intestinal inflammation, we examined GPR4 mRNA expression in WT-DSS and WT-control mice by real-time PCR. Our results demonstrated that GPR4 mRNA expression was upregulated in the inflamed colonic tissue of WT-DSS mice by nearly 2.7 fold when compared to normal controls (Fig. 4A). No GPR4 mRNA expression was detected in colon tissues of GPR4 KO mice, confirming the deficiency of GPR4 in the KO mice (data not shown). Furthermore, we measured the GPR4 mRNA expression in IBD and normal human intestinal tissues by real-time PCR. We used a cDNA array containing 7 normal colon, 26 active colitis, and 14 Crohn’s tissue samples. We observed a ~4.7 fold increase of GPR4 mRNA expression in human IBD lesions compared to normal human intestinal tissues (Fig. 4B). These data collectively demonstrate that GPR4 expression is increased in IBD intestinal lesions and could be involved in the pathogenesis of the disease.

**Figure 4.**
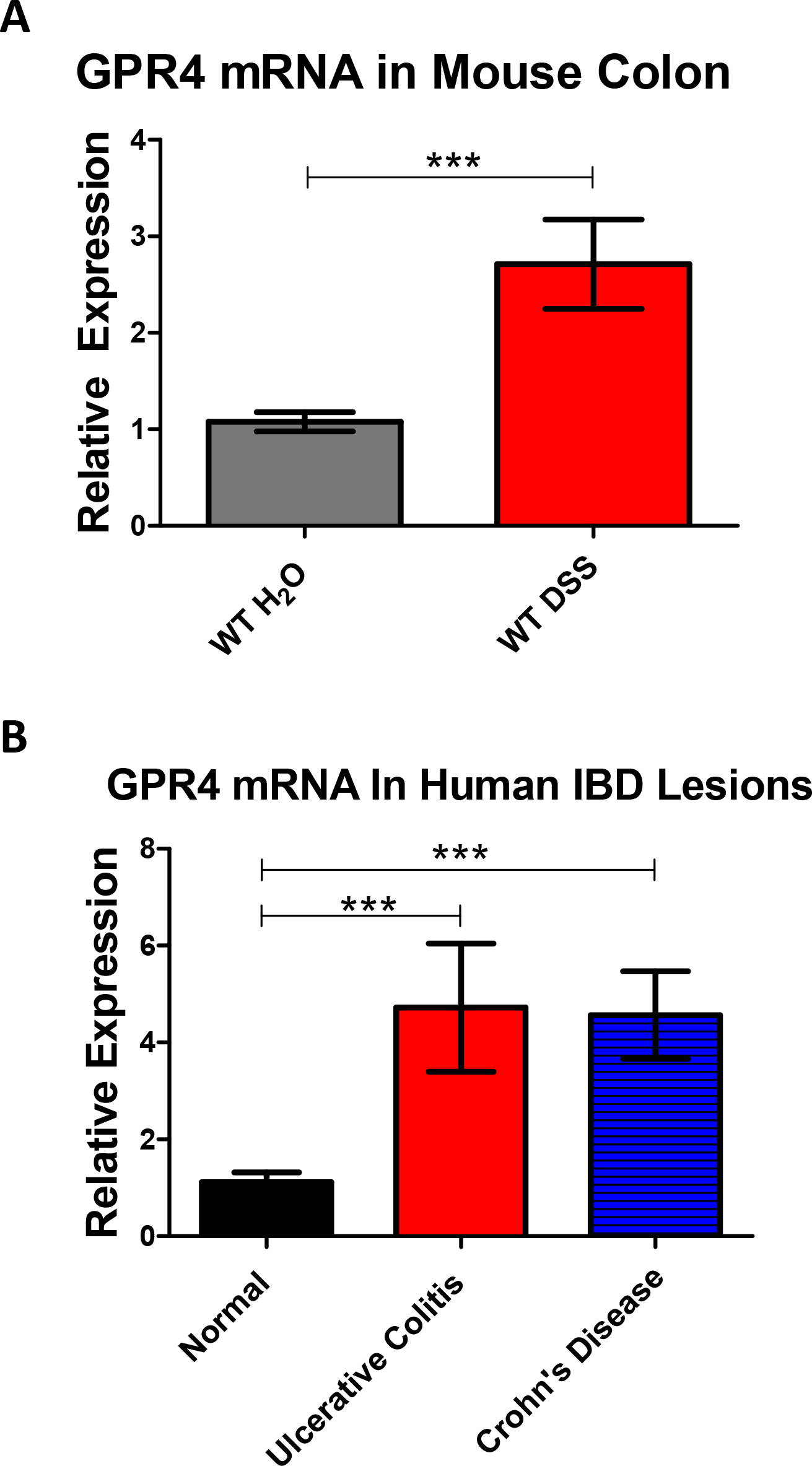
| GPR4 mRNA expression in human and mouse IBD tissues. Expression levels of GPR4 in diseased and non-diseased intestinal tissues were assessed. GPR4 mRNA was increased in IBD lesions of human and mouse intestinal tissues when compared to normal intestinal tissues. GPR4 mRNA expression in mouse WT-DSS colonic tissue compared to WT-control tissues (A). GPR4 mRNA expression in human normal, ulcerative colitis, and Crohn’s intestinal tissues (B). Data are presented as mean ± SEM (*** P< 0.001).

### GPR4 expression is primarily localized in vascular endothelial cells in intestinal tissues

In order to characterize the expression pattern of GPR4 in the mouse colon and cecum, we performed immunohistochemistry for green fluorescent protein (GFP) as a surrogate marker for GPR4 due to the lack of an antibody that can reliably detect endogenous GPR4 protein. GPR4-deficient mice were generated by replacing the GPR4 coding region with an internal ribosome entry site (IRES)-GFP cassette under the control of the GPR4 gene promoter as previously described ^9^. Therefore, GFP expression in mouse tissues serves as a surrogate marker for endogenous GPR4 expression. GFP expression was predominately detected in the endothelial cells (ECs) of blood vessels, including arteries, veins, and microvessels of the cecum and colon in GPR4 KO-control mice (Fig.5A-E, SI:5A-B). Additionally, GFP expression could also be observed in the specialized high endothelial venules (HEVs) in mesenteric lymph nodes and microvessels adjacent to ILFs (Fig. 5A-F). GFP expression could not, however, be significantly detected in lymphatic endothelial cells (Fig. 5A). In addition to GFP expression in ECs, GFP expression could be detected in histiocytes (macrophages) located in the sinus of the mesenteric lymph nodes (Fig. 5F). No GFP signal could be detected in WT untreated control intestinal tissues, with the exception of background signal on the epithelium, luminal content, and adipose tissue (SI:3, SI:5E-F). The GFP expression pattern is in accordance with previously published results showing that GPR4 is expressed in several types of cultured vascular endothelial cells and isolated monocytes/macrophages ^18, 50 –52^. The role of GPR4 in macrophages is currently unknown and further studies will need to be conducted in the future to elucidate the functional and molecular role of GPR4 in macrophages.

**Figure 5.**
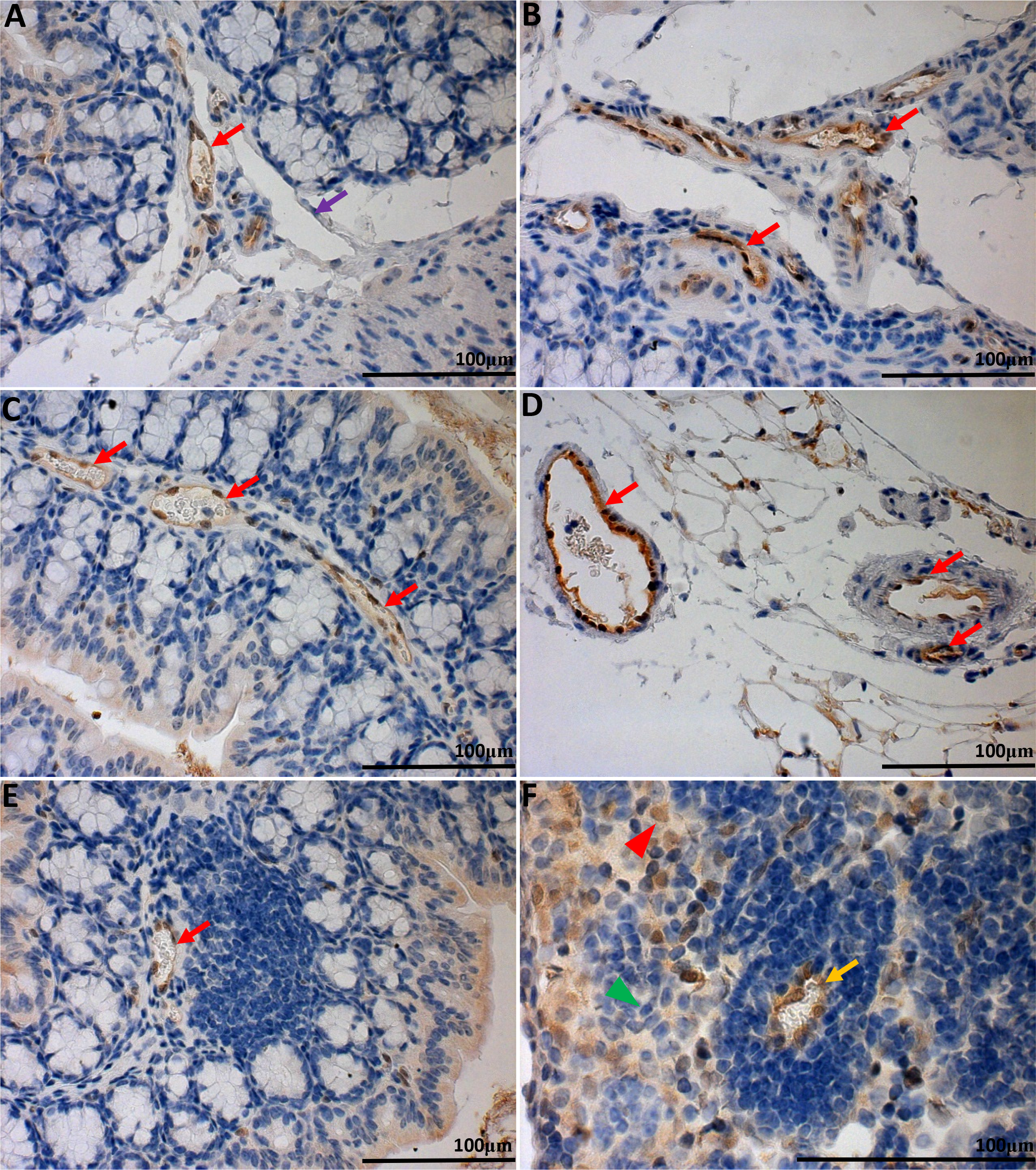
| GFP knock-in as a surrogate marker for GPR4 expression in GPR4 KO control mouse colon and lymph tissues. To assess the localization of GPR4 in intestinal tissues, we performed IHC of GFP in intestinal and lymph tissues. GFP expression was visualized as brown signals in the intestinal microvascular endothelial cells, ex-mural blood vessels, and mesenteric lymph node high endothelial venules (HEVs). GFP expression was not observed in lymphatic ECs. Colonic GPR4 KO-control mouse blood vessel, artery, and lymphatic vessel (A&B). Transverse fold microvessels (C); ex-mural blood vessel and arteries (D). Microvessels adjacent to isolated lymphoid follicles (E); mesenteric lymph node HEVs, and histiocytes (F). No GFP signal detected in WT untreated control tissues (SI:5). 40× and 63× microscope objectives. Red arrow heads indicate histiocytes (macrophages) and green arrow heads indicates lymphocytes. Red arrows indicate blood vessels, yellow arrows indicate HEVs, and purple arrows indicate lymphatics.

Upon examination of GFP expression in the inflamed colon and cecal tissues of GPR4 KO-DSS mice, similar localization of GFP in the endothelial cells of arteries, veins, microvessels, and HEVs could be observed as the GPR4 KO control mice (Fig. 6A-E, SI:5C-D). GFP expression could also be detected in mucosal macrophages in inflamed lesions of the GPR4 KO-DSS mice as well as the sinus regions of the mesenteric lymph nodes (Fig. 6B, F). In accordance with the WT untreated control tissues, no GFP signal could be detected in WT-DSS tissues with the exception of minor background staining of the luminal epithelium, luminal content, and adipose tissue (SI:4, SI:5G-H).

**Figure 6.**
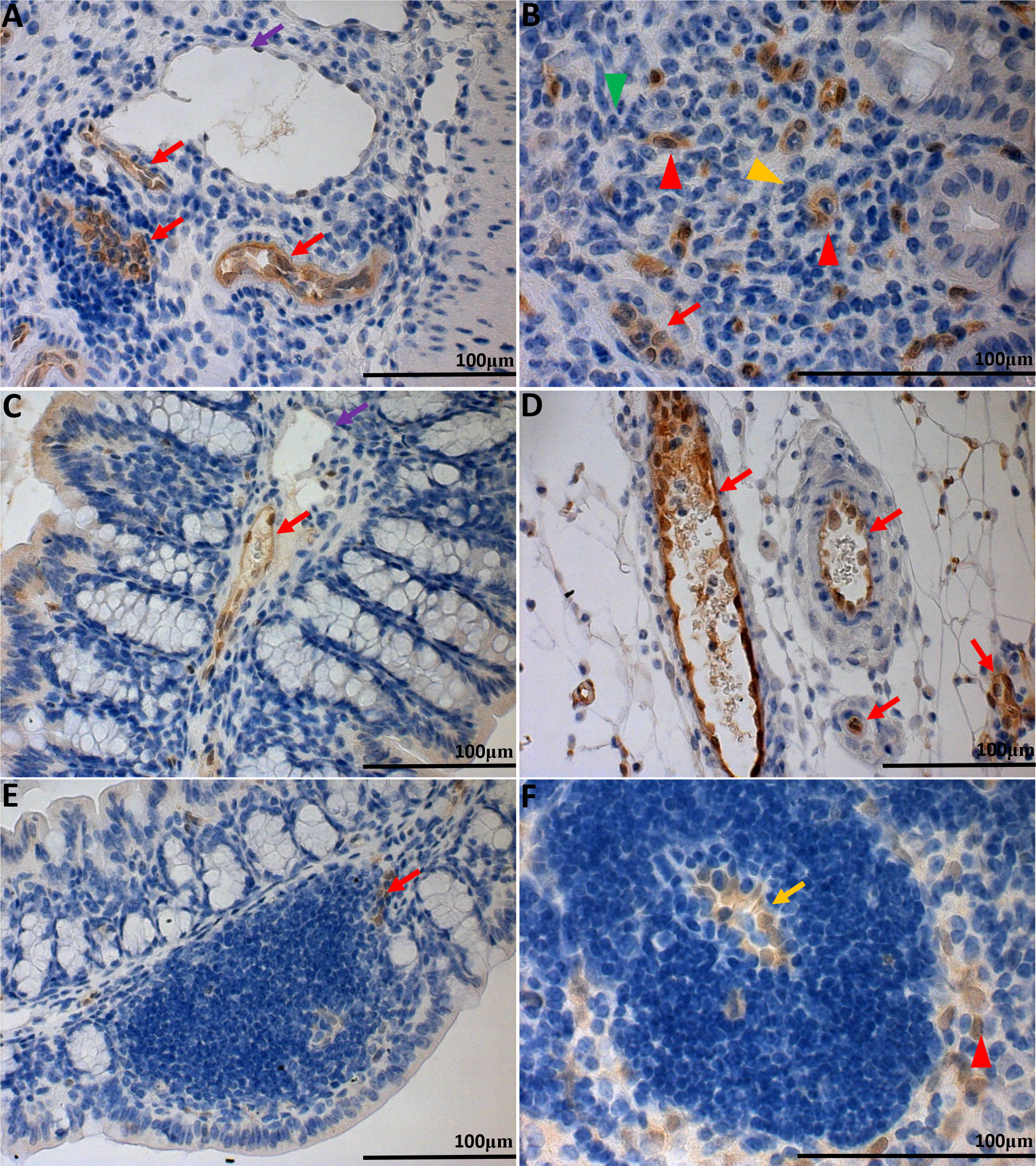
| GFP knock-in as a surrogate marker for GPR4 expression in GPR4 KO-DSS mouse colon and lymph tissues. To examine the expression of GPR4 in inflamed intestinal tissues, we performed IHC of GFP in GPR4 KO-DSS tissues. GFP expression could be visualized as brown signals in the intestinal microvascular endothelial cells, ex-mural blood vessels, mesenteric lymph node HEVs, and macrophages. No expression of GFP could be detected in lymphatic ECs. Colonic GPR4 KO-DSS blood vessel, artery, and lymphatic (A), resident macrophages in inflamed lesions (B), transverse fold ECs (C), ex-mural blood vessels (D), isolated lymphoid follicle vessels (E), and mesenteric lymph node HEVs (F). No expression of GFP could be detected in WT-DSS control tissues (SI:6). 40× and 63× microscope objectives. Red arrow heads indicate macrophages, yellow arrow heads indicate neutrophils, and green arrow heads indicates lymphocytes. Red arrows indicate blood vessels, yellow arrows indicate HEVs, and purple arrows indicate lymphatics.

Due to the localization of GPR4 in HEVs traversing lymphoid tissues such as mesenteric lymph nodes and in microvessels adjacent to ILFs in the mucosa, GPR4 could regulate lymphoid tissue expansion in a manner consistent with previous publications demonstrating GPR4 in ECs increases numerous cytokines, chemokines, and adhesion molecules regulating leukocytes interaction with ECs ^17, 18^. GPR4 could be involved in increasing the passage of leukocytes critical for inflammatory responses in secondary and tertiary lymphoid tissues as is observed in GPR4 KO-DSS mice having reduced mesenteric lymph node volume (Fig. 1D) and ILF development (Fig. 3, SI:2).

### GPR4 deficiency decreases the expression of inflammatory genes in the colon ofDSS-treated mice

As previous GPR4 inhibitor and shRNA knockdown studies have shown that inhibition of GPR4 reduces the expression of adhesion molecules and numerous inflammatory genes in endothelial cell cultures ^17, 18^, we sought to examine a selection of inflammatory genes expressed in WT and GPR4 KO whole colon tissue. Given the diverse cell population in the inflamed colon, coupled with the focal nature of IBD, inflammatory molecules such as VCAM-1, E-selectin and ICAM-1 can be expressed by a variety of stromal and immune cells in addition to endothelial cells ^53–55^. As GPR4 is expressed primarily in vascular endothelial cells, much of the inflammatory molecule expression in other cell types within the inflamed colon tissue is not regulated by GPR4. Therefore, the effects of EC regulation by GPR4 can be masked by gene analysis of whole tissue colon segments. In spite of this limitation, using real-time PCR analysis of whole colon tissue segments, we observed the mRNA expression of adhesion molecules E-selectin, ICAM-1, VCAM-1, and MAdCAM-1 were reduced, although not statistically significant with this sample size, in the GPR4 KO-DSS mouse colon compared to the WT-DSS colon (Fig. 7A-D). In addition to adhesion molecule expression, mRNA levels of COX-2 were reduced in GPR4 KO-DSS mice whereas CXCL2 showed little difference between WT-DSS and GPR4 KO-DSS mice (Fig. 7E-F).

**Figure 7.**
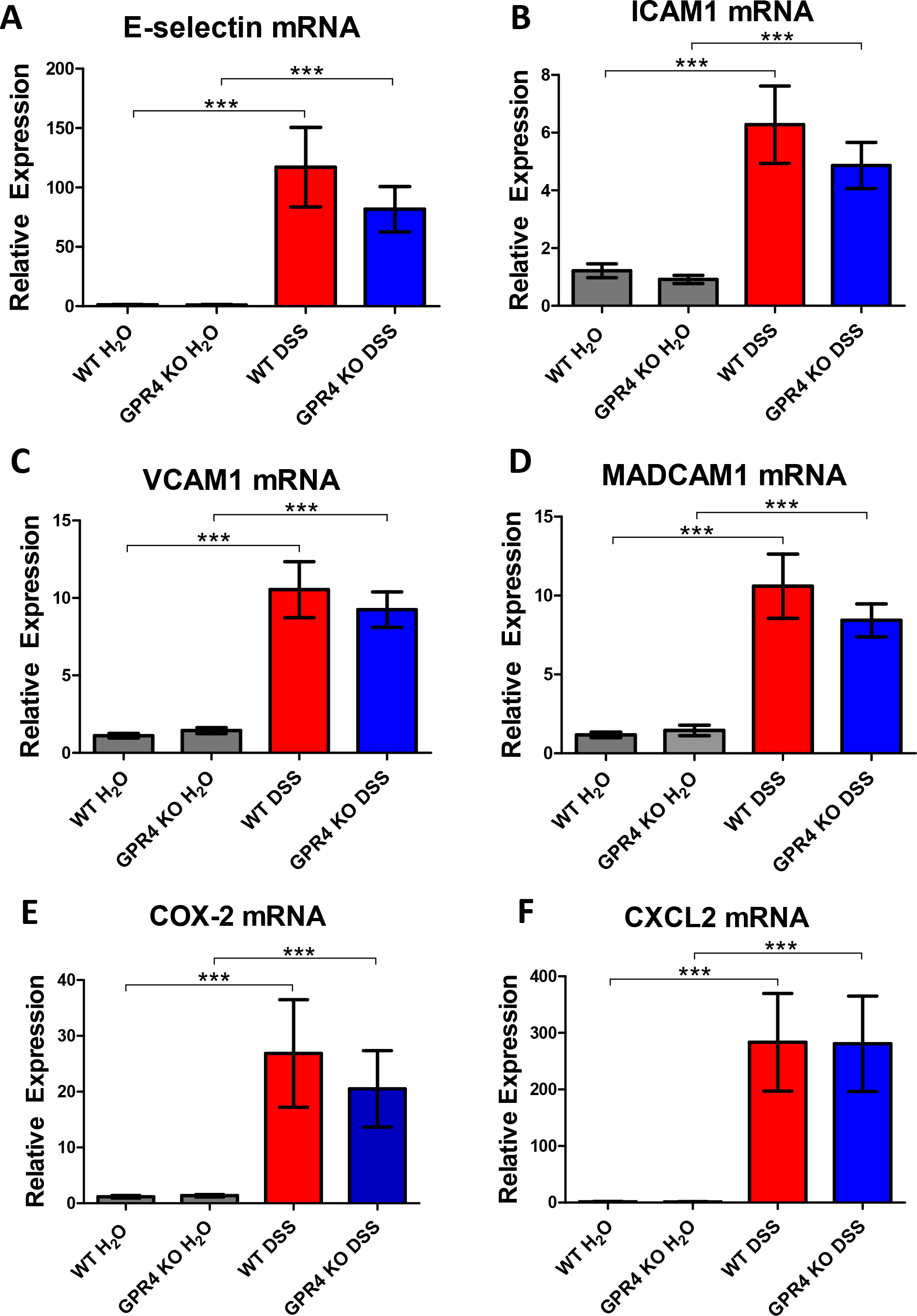
| Real-time PCR analysis of inflammatory gene expression. Inflammatory gene expression was evaluated in whole colon tissue segments to assess the contribution of GPR4, among other cells not regulated by GPR4, to inflammatory molecule expression. The expression of the inflammatory genes was significantly increased in both WT and GPR4 KO DSS colon tissues when compared to control colon tissues. GPR4 KO-DSS mice exhibited a trend of reduced pro-inflammatory gene expression when compared to WT-DSS mice. Adhesion molecules E-selectin (A), ICAM-1 (B), VCAM-1 (C), and MAdCAM-1 (D) were analyzed along with inflammatory enzyme COX-2 (E), and chemokine CXCL2 (F). Data are presented as mean ± SEM (*** P< 0.001).

Furthermore, correlating GPR4 mRNA expression in WT-DSS colonic tissues to inflammatory gene expression, we were able to see a statistically significant positive correlation between increased GPR4 mRNA expression and inflammatory gene expression such as E-selectin and COX-2 (Fig. 8A, E). With the exception of CXCL2; ICAM-1, VCAM-1, and MAdCAM-1 showed a strong trend in correlation with GPR4 mRNA expression that has not yet reached statistical significance given the current sample size (Fig. 8B, C, D, F).

**Figure 8.**
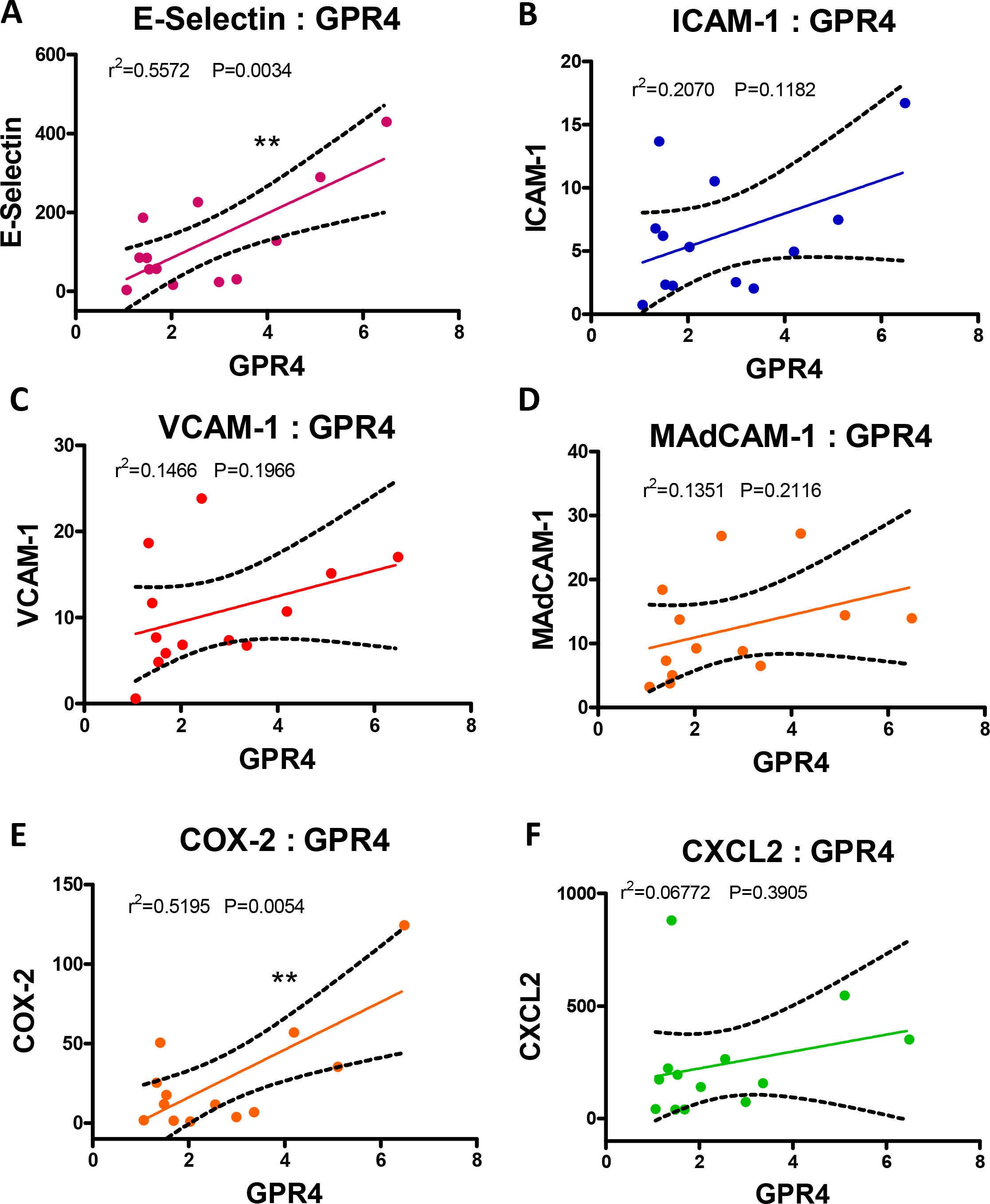
| Correlation between GPR4 mRNA expression and inflammatory gene expression. To further characterize GPR4 regulated inflammatory gene expression, GPR4 mRNA expression was correlated with inflammatory gene expression from WT-DSS colon segments (n=13). Each dot represents the data from an individual mouse. GPR4 mRNA expression positively correlates with increased inflammatory gene expression when analyzing (A) E-selectin, (B) ICAM-1, (C) VCAM-1, (D) MAdCAM-1, (E) COX-2, and (F) CXCL2. *(**P* < 0.01)

To further examine the effects of GPR4 regulated inflammatory molecule expression at the cellular level, we performed immunohistochemistry to examine the expression of E-selectin and VCAM-1 in the vascular endothelium. Overall, the E-selectin and VCAM-1 protein expression was increased in the DSS-treated inflamed WT and GPR4 KO mouse intestinal tissues when compared to the non-treated control tissues. Immunohistochemical analysis of E-selectin revealed its expression in the vascular endothelium and some colon epithelial cells. One report confirms the expression of E-selectin in colon epithelial cells ^54^. We observed ECs in GPR4 KO-DSS mouse colons and cecums had reduced E-selectin expression when compared to WT-DSS mice (Fig. 9A-D, 10A-D). Immunohistochemical analysis of VCAM-1 revealed its expression on a variety of cell types in addition to the ECs within the inflamed colon and cecum. Expression could be observed in the mucosa, submucosa, and muscularis externa. Overall, total VCAM-1 expression was visibly reduced and less extensive in the GPR4 KO-DSS mice compared to WT-DSS mice, and an appreciable reduction in VCAM-1 signal could be discerned in the mucosal endothelial cells themselves within the colon and cecum tissues (Fig. 9E-H, 10E-H).

**Figure 9.**
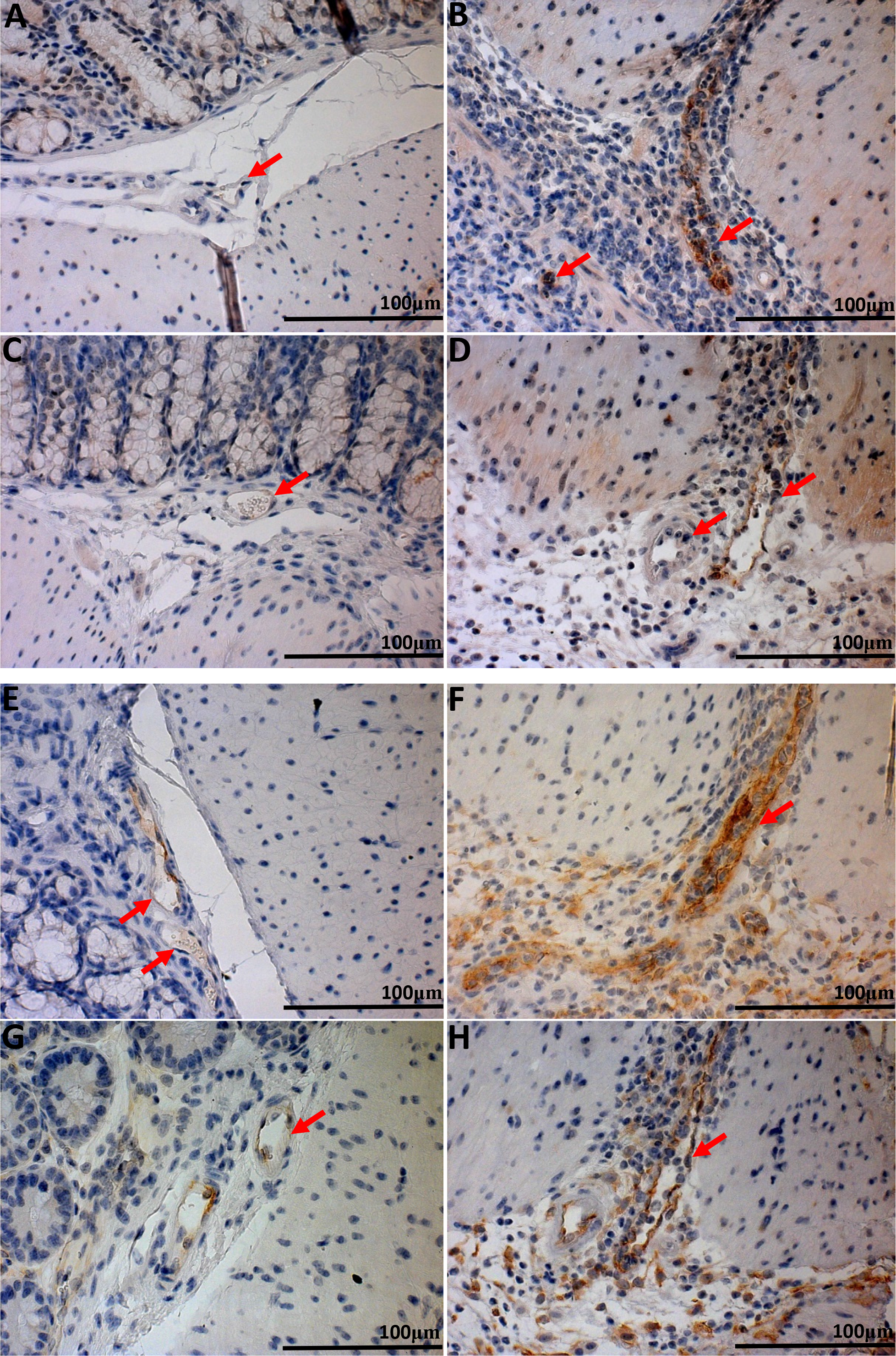
| Immunohistochemical analysis of E-selectin and VCAM-1, respectively in mouse colon tissues. As whole tissues are not ideal for analyzing endothelial cell specific gene expression, we performed IHC to analyze adhesion molecules E-selectin and VCAM-1 expression in ECs within the tissue. GPR4 KO-DSS mice have reduced E-selectin and VCAM-1 expression in colonic mucosal vasculature when compared to WT-DSS mice. E-selectin expression could be visualized as brown signals in WT-control (A), WT-DSS (B), GPR4 KO-control (C), and GPR4 KO-DSS (D) colon tissues. VCAM-1 expression could be visualized in WT-control (E), WT-DSS (F), GPR4 KO-control (G), and GPR4 KO-DSS (H) colon tissues. 40× microscope objective. Red Arrows indicate blood vessels.

**Figure 10.**
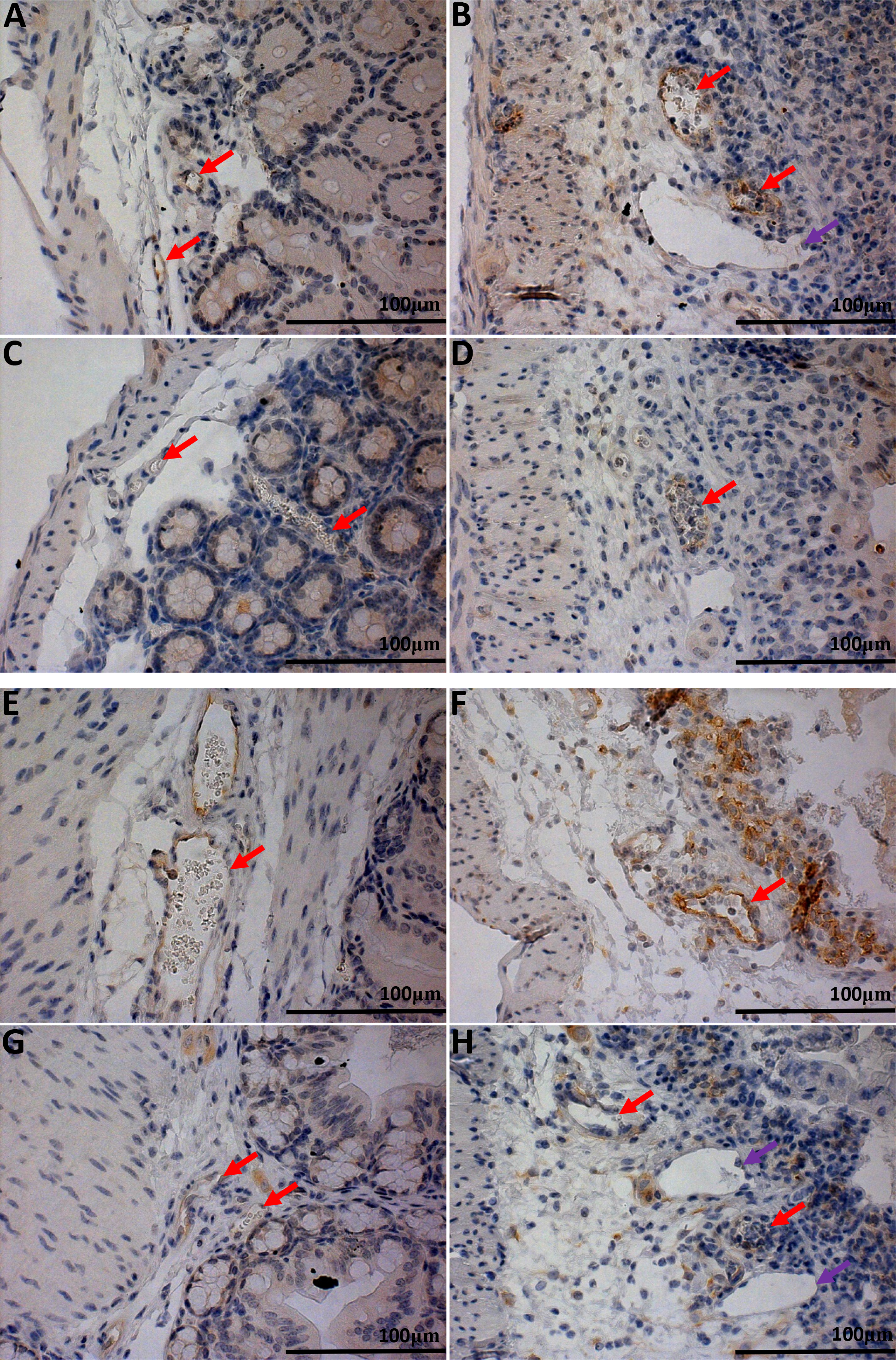
| Immunohistochemical analysis of cecum adhesion molecule expression in ECs. In addition to the colon, cecum tissues were examined for adhesion molecule expression between WT and GPR4 KO mice. Similar to colon, GPR4 KO-DSS mice had a reduction in the expression of E-selectin and VCAM-1 in ECs. E-selectin expression could be visualized as brown signals in WT-control (A), WT-DSS (B), GPR4 KO-control (C), and GPR4 KO-DSS mucosal blood vessels (D). VCAM-1 expression could be visualized in WT-control (E), WT-DSS (F), GPR4 KO-control (G), and GPR4 KO-DSS mucosal blood vessels (H). 40× microscope objective.

Taken together, GPR4 appears to increase the expression of E-selectin, ICAM-1, VCAM-1, and MAdCAM-1 in the inflamed colon as real-time PCR results showed a reduction in inflammatory molecule expression in GPR4 KO-DSS mice compared to WT-DSS mice from whole tissue lysate. Furthermore, directly examining the expression of E-selectin and VCAM-1 in colonic and cecal blood vessels confirmed the reduction of adhesion molecules expressed in ECs of GPR4 KO-DSS mice compared to WT-DSS mice. These data suggest GPR4 could increase intestinal inflammation through the regulation of endothelial inflammation and leukocyte infiltration into inflamed gastrointestinal tissues.

## Discussion

This study demonstrates that GPR4 deficiency alleviates intestinal inflammation in the DSS-induced acute IBD mouse model, suggesting GPR4 is a pro-inflammatory gene in IBD. These results corroborate previous studies showing GPR4 contributes to the inflammatory response in endothelial cells by isocapnic and hypercapnic acidotic stress *in vitro* ^17, 18^. GPR4 significantly increased endothelial cell and leukocyte interaction through the up-regulation of cytokines, chemokines, and adhesion molecule gene expression ^17–19^. Our *in vivo* data described in this report provide novel insights into the vascular inflammatory response in intestinal inflammation. In keeping with previous results describing the role of GPR4 in various endothelial cell types ^17–19^, our results suggest that GPR4 increases the expression of cytokines, chemokines, and adhesion molecules in intestinal endothelial cells and potentiates intestinal inflammation by increasing leukocyte recruitment to the gut and extravasation from blood to intestinal mucosa in response to acidic tissue microenvironments.

Acid sensing is critical for cells in the maintenance of proper cellular functions. When improper pH homeostasis occurs, various pathological conditions can arise. Extracellular tissue pH is tightly regulated around 7.4, while intracellular pH is slightly more acidic (pH 7.2). If cells are unable to maintain this narrow pH range, cell death will occur ^56, 57^. Cells must be able to sense and modulate their processes in response to an altered extracellular pH gradient. pH sensing can occur through acid sensing ion channels (ASICs), transient receptor potential (TRP) channels, and the GPR4 family of proton-sensing GPCRs ^4, 58^. Recently, GPR4 has been implicated in regulating systemic pH through the renal system ^59^. GPR4 is expressed in the kidney and involved in the pH sensing function of kidney collecting duct cells and regulating renal acid secretion in response to systemic acidotic challenges. In addition to the renal system, GPR4 expression in retrotrapezoid nucleus (RTN) neurons mediate the central respiratory chemoreflexes to altered systemic CO_2_/pH homeostasis ^60^. This study showed GPR4 is required for RTN neurons to respond to elevated brain pCO_2_. GPR4 is also involved in local pH responses where tissue becomes acidic due to pathological conditions such as ischemia, hypoxia, and inflammation ^1–3^. Indeed, tissue acidosis is a hallmark of inflammation.

Numerous studies have demonstrated that acute and chronically inflamed tissues are characterized by acidosis and can modulate the function of both immune cells and stromal cells ^22, 23^. The reduced tissue pH is owing to multiple factors in the context of inflammation. Increased leukocyte infiltrates quickly deplete available O_2_ and cells in the hypoxic tissue switch from aerobic to anaerobic glycolysis generating lactic acid. Neutrophils are typically the first responders to an inflammatory stimuli which is often due to bacterial infiltration. These bacteria can acidify the inflammatory microenvironment owing to accumulation of short chain fatty acids as microbial metabolic by-products ^61, 62^. Neutrophils and macrophages can attempt to eliminate these harmful bacteria through respiratory bursts, which can further acidify the microenvironment ^63, 64^. Interestingly, reports find that the colonic lumen of patients with active UC is more acidic than the normal colon ^65, 66^. There are some varying reports regarding intraluminal colonic pH ^67^, however, the consensus seems to hold the lumen of active UC patients are more acidic than control patients ^28, 44^.

Using the GFP knock-in as a surrogate marker for GPR4 expression in GPR4 KO intestinal tissues, we were able to observe GFP expression on ECs of blood vessels in the muscularis externa, mucosa, and transverse folds (Fig. 5, 6, SI:5). Interestingly, however, no expression of GFP was visible in lymphatic ECs. Increased density of GFP-positive blood vessels could clearly be observed in the context of inflamed intestinal lesions of GPR4 KO-DSS mice where tissue acidosis is co-existent with inflammation. It is possible that the increased GPR4 expression could be correlated with increased blood vessel density in the inflamed tissues. Increased angiogenesis is common in the mucosa of IBD patients and these blood vessels grow into inflamed, acidic tissues. With previous reports showing GPR4 expression is increased by TNF-α and H_2_O_2_, of which is commonly expressed in the milieu of the inflamed gut mucosa ^15^, GPR4 expression could be upregulated in vascular endothelial cells and potentiate inflammation in response to the acidic microenvironment. Consistently, the increased mRNA expression of GPR4 by nearly 2.7 fold was observed in WT-DSS mice and 4.7 fold in human intestinal IBD lesions (Fig. 4). Due to the lack of a reliable GPR4 antibody, we cannot directly examine the expression of GPR4 protein. Using IHC methods for detecting the knock-in GFP expression can serve as a surrogate marker to localize GPR4 expression in intestinal tissues (Fig. 5, 6, SI:5), but does not necessarily correlate to the level of GPR4 protein and cannot be used to quantitatively measure GPR4 protein expression between non-inflamed and inflamed ECs. Nonetheless, increased GPR4 expression correlates with the observed IBD pathology and can contribute to the pathogenesis of the disease.

High endothelial venules (HEVs) are specialized ECs for lymphoid tissues. In addition to visualizing GFP expression in mucosal vasculature, we were also able to observe GFP expression in HEVs of mesenteric lymph nodes and microvessels of ILFs (Fig. 5, 6). GPR4 could play a substantial role in the expansion of secondary and tertiary lymphoid tissue. Based on macroscopic disease indicators, we observed the average volume of mesenteric lymph nodes from GPR4 KO-DSS mice were similar in size to WT control mice and ~40% reduced when compared to WT-DSS mice (Fig. 1). Additionally, a significant reduction can be observed in isolated lymphoid follicle numbers between WT-DSS and GPR4 KO-DSS (Fig. 3). GPR4 could play a novel role in regulating the passage of leukocytes, such as lymphocytes and antigen presenting cells into lymph tissue and thereby regulate the adaptive immune response.

As immunologic, environmental, and genetic factors have been clearly shown to contribute to the development and progression of IBD, so too are these pH-sensing GPCRs implicated in the regulation of IBD within these categories. Ovarian cancer G protein-coupled receptor 1 (OGR1) has recently been implicated as a regulator of intestinal inflammation through the control of macrophage inflammatory responses ^43^. Using IL-10^−/−^ (knockout) mice for the development of chronic spontaneous intestinal inflammation, OGR1 deficiency alleviated mucosal inflammation when compared to control mice. This group demonstrates OGR1 expression, among other pH-sensors, is exclusively upregulated by TNF-α, PMA, and LPS in macrophages and potentiates intestinal inflammation. Interestingly, using IHC analysis, we observed GFP (a surrogate marker for GPR4) expression in mouse macrophages in both the lymph node sinus region and inflamed mucosal lesions of GPR4-KO mice (Fig. 5, 6). This observation is consistent with previous results showing that GPR4 is expressed in purified monocytes and macrophages ^50–52^. However, the role of GPR4 in macrophages is unknown and further studies must be done to characterize the functional role of GPR4 in macrophages. This endeavor may be confounded by genetic redundancy as the whole family of proton sensing GPCRs are expressed in macrophages. Additionally, in keeping with these observations that GPR4 KO and OGR1 KO mice have a similar phenotype under intestinal inflammation, it is possible GPR4 and OGR1 may have redundant roles. Additional studies using OGR1 and GPR4 double knockouts will need to be done to reveal further biological functions of these receptors. In addition to ORG1 regulation of inflammatory cells in the gut, the receptor G2A (G2 accumulation) is expressed on leukocytes and is potentially involved in host-microbial gut interactions. A commensal microbial metabolite called commendamide isolated from the gut bacterium *Bacteroides* could activate G2A and was hypothesized to provide an antiinflammatory role in IBD ^68^. Finally, T cell death associated gene 8 (TDAG8) has recently been presented as an IBD susceptibility candidate gene ^69^. Polymorphisms in the *TDAG8* (*GPR65*) gene have been linked with increased risk of developing IBD based on several genome-wide association study (GWAS) efforts. A recent study showed that TDAG8 (GPR65) deficient mice have increased susceptibility to bacteria-induced colitis ^70^. Collectively, the GPR4 family of receptors are emerging as regulators of intestinal inflammation.

Here we propose a mechanism of how GPR4 specifically contributes to intestinal inflammation (Fig. 11). As mentioned earlier, we have previously reported that GPR4 can stimulate the expression of adhesion molecules, chemokines, and other inflammatory genes in a variety of endothelial cells in response to isocapnic and hypercapnic acidosis ^17, 18^. Notably, some of these inflammatory molecules are E-selectin, VCAM-1, and ICAM-1. These adhesion molecules are involved in the tethering and firm adhesion of leukocytes to endothelial cells and are regulated by GPR4. To confirm a functional role for the GPR4-dependent expression of adhesion molecules, we performed adhesion assays under static and flow conditions. We observed an increase in leukocyte adhesion to endothelial cells in a GPR4 dependent manner ^17, 18^. The leukocyte adhesion was reduced when treated with a GPR4 inhibitor in a dose dependent manner or by GPR4 shRNA knockdown in ECs.

**Figure 11.**
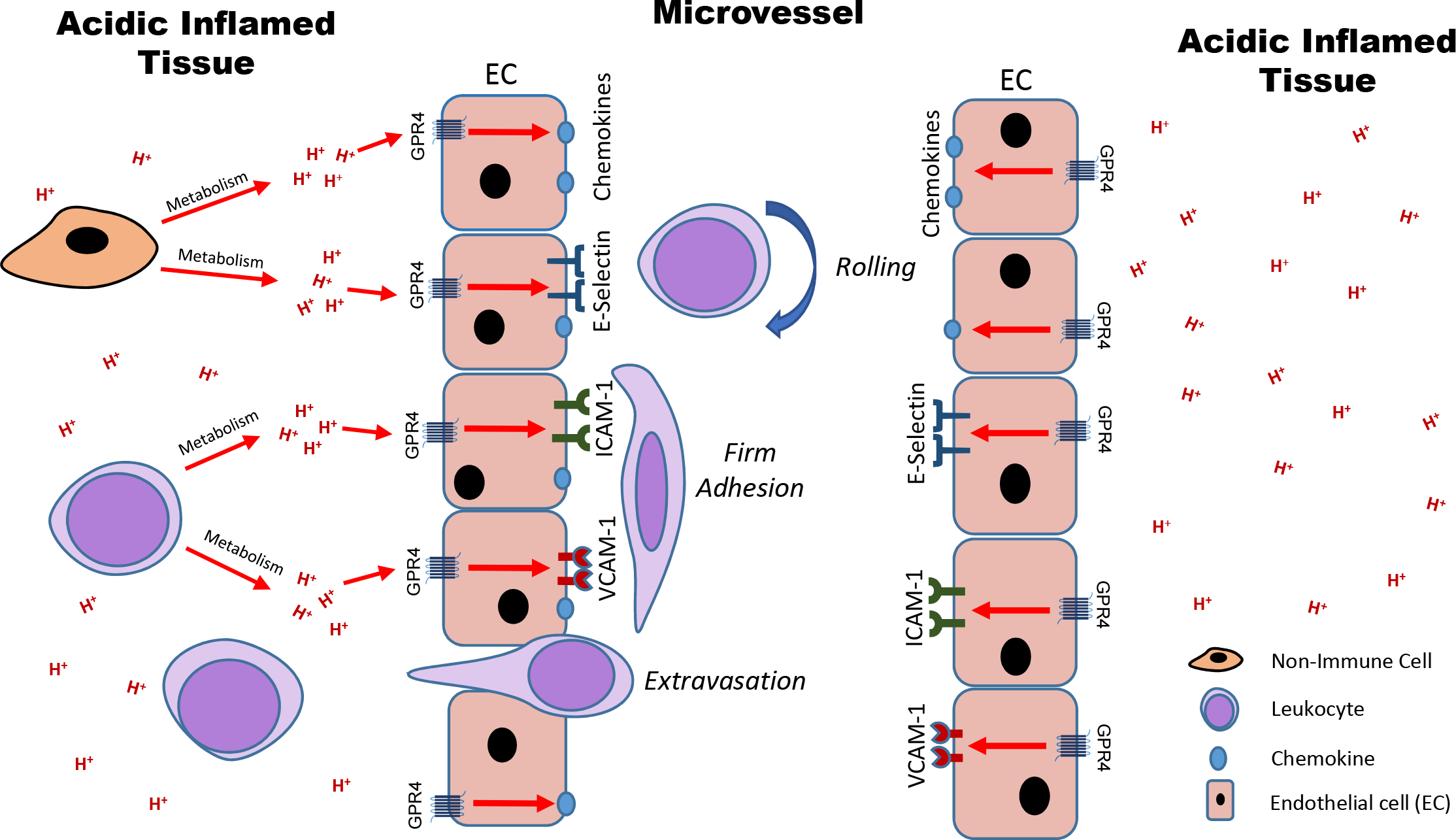
| Model of proposed mechanism for the governance of endothelial cell inflammatory responses by GPR4. GPR4 can be activated by protons in the acidic microenvironment and increase the expression of adhesion molecules (e.g. E-selectin, ICAM-1, and VCAM-1) and chemokines (e.g. CCL20, CXCL2, and IL-8) for the recruitment and adherence of leukocytes to the endothelium. Increased leukocyte extravasation will occur into the inflamed tissue and potentiate local inflammation.

Consistent with our previous publications on the role of GPR4 in cellular adhesion, we observed GPR4 increases pro-inflammatory molecule mRNA expression in WT mice (Fig. 7). Even though whole colon tissue is not ideal for the examination of EC specific gene expression, we were able to observe a trend of reduction in E-selectin, VCAM-1, ICAM-1, MAdCAM-1, and COX-2 mRNA in GPR4 KO-DSS mice compared to WT-DSS mice. Furthermore, inflammatory gene expression positively correlated with GPR4 mRNA gene expression in WT-DSS colon tissues (Fig. 8), suggesting the inflammatory genes are co-regulated by GPR4. Using immunohistochemistry, we were able to observe the reduction in E-selectin and VCAM-1 specifically in the blood vessels of GPR4 KO-DSS mice compared to WT-DSS mice (Fig. 9, 10). These observations are consistent with our previous reports showing GPR4 increases the expression of adhesion molecules, chemokines and other inflammatory genes ^17, 18^. Additionally, histopathological analysis confirmed a reduction of leukocyte infiltration into the mucosa of GPR4-KO-DSS mice compared to WT-DSS mice (Fig. 2), suggesting GPR4 deficiency can reduce leukocyte infiltration into inflamed tissue through the regulation of adhesion molecules in ECs *in vivo.* We propose the reduced IBD clinical severity, pro-inflammatory molecule expression, and isolated lymphoid follicle development in GPR4-KO-DSS mice is owing to the GPR4 dependent effects on pro-inflammatory molecule expression, leukocyte trafficking, and extravasation into inflamed mucosal tissue in the gut (Fig. 11). While focused on IBD in this study, a similar GPR4 regulating mechanism can potentially be extrapolated to many other inflammatory disorders.

Inhibition of leukocyte-endothelial cell interactions are an attractive approach to treat inflammatory diseases as the inhibition of leukocyte adhesion would reduce the influx of inflammatory cells into inflamed tissue. Currently, FDA approved therapeutics targeting EC-leukocyte interaction, such as vedolizumab and natalizumab, are used for IBD treatment ^31, 32^. Even though these drugs have been shown to reduce aberrant inflammatory responses, the efficacy of targeting specific inflammatory mediators can be reduced over time through host immune compensatory mechanisms. Examples can be observed in the recently failed efforts to attenuate intestinal inflammation by supplementing interleukin-10 in IBD patients ^71^. GPR4 is a strong potential candidate for targeted therapy, as it is upstream of the predominate EC-leukocyte targets currently used in IBD therapy and can up-regulate adhesion molecules and chemokines involved in both tethering and firm adhesion of leukocytes to the endothelium as well as leukocyte activation. Use of GPR4 inhibitors could prove as a valuable tool to inhibit inflammation by reducing leukocyte recruitment and adhesion to inflamed tissues. Recently, our group and others have demonstrated the effectiveness of the GPR4 inhibitors in the selective targeting of GPR4 and inhibition of GPR4 target gene expression ^17, 19, 72^ Additionally, GPR4 antagonists have been used *in vivo* and no obvious toxicities have been reported ^20, 72^. A recent study showed that GPR4 antagonists provided therapeutic benefits in a myocardial infarction mouse model ^72^. Use of GPR4 antagonists could prove as a novel therapeutic in the treatment of IBD and other inflammatory diseases.

## Acknowledgement

We would like to thank Connor Hart and Lixue Dong for their excellent technical assistance. Additionally, we would like to thank Dr. Owen N. Witte for providing the GPR4 knockout mouse strain and Dr. Mary J. Thomassen’s lab for generously sharing equipment and reagents. This study was supported in part by Brody Brothers Endowment Fund, American Heart Association, and Vidant Cancer Research and Education Fund (to L.V.Y).

## Author contributions

EJS, KL, and LVY designed the experiments; EJS, NRL performed the majority of the experiments and KL, JZO, CRJ, EAK, DO, LVY performed some experiments; QC, HH, VB, and JGF performed histopathological analyses; EJS, NRL, and LVY analyzed data; EJS and LVY wrote the manuscript; all authors reviewed and edited the manuscript.

## Supplementary Information

**SI: 1.**
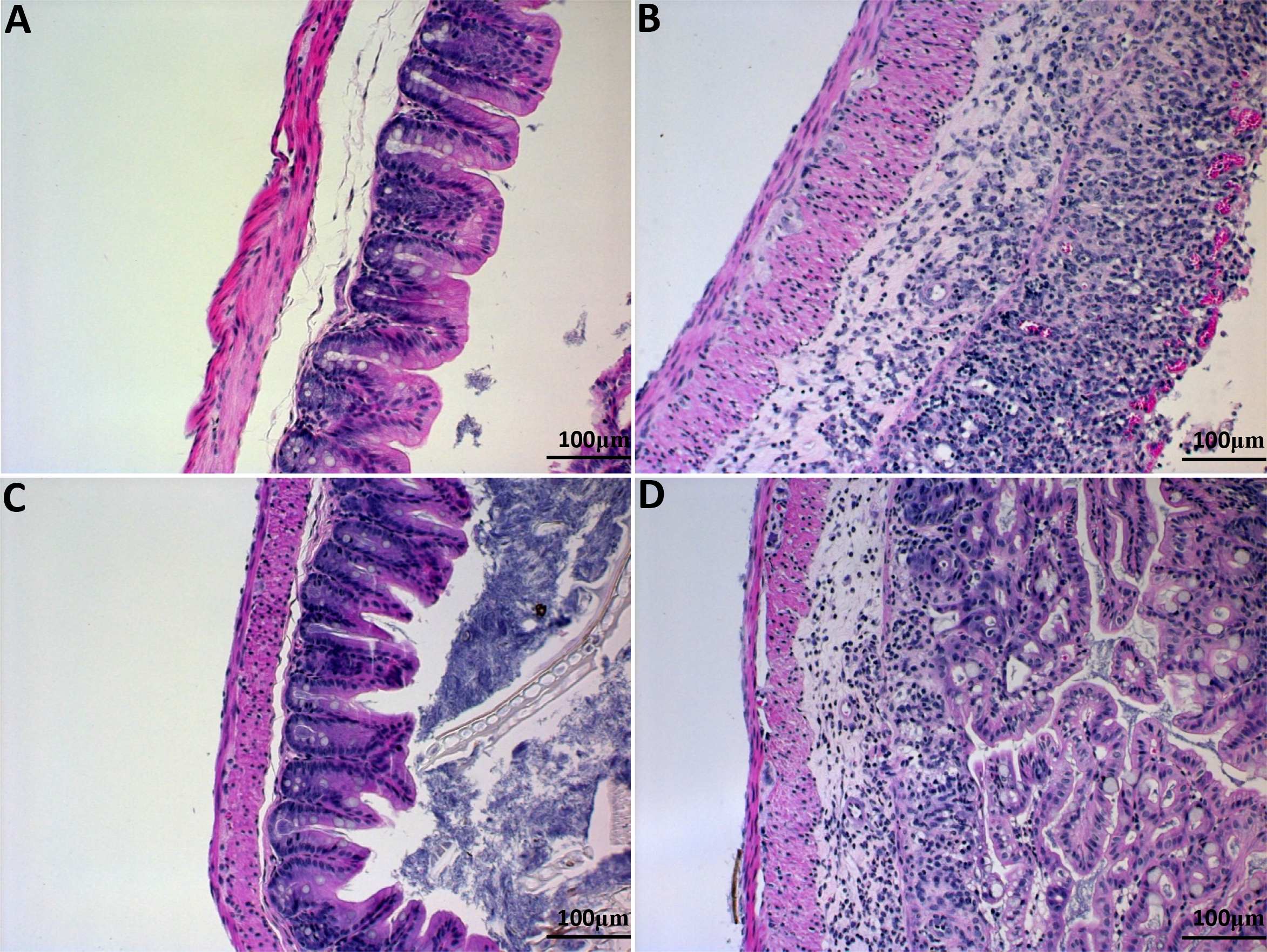
| H&E representative pictures of cecum in WT control (A), WT DSS (B), GPR4 KO control (C), and GPR4 KO DSS (D). 20× microscope objective.

**SI: 2.**
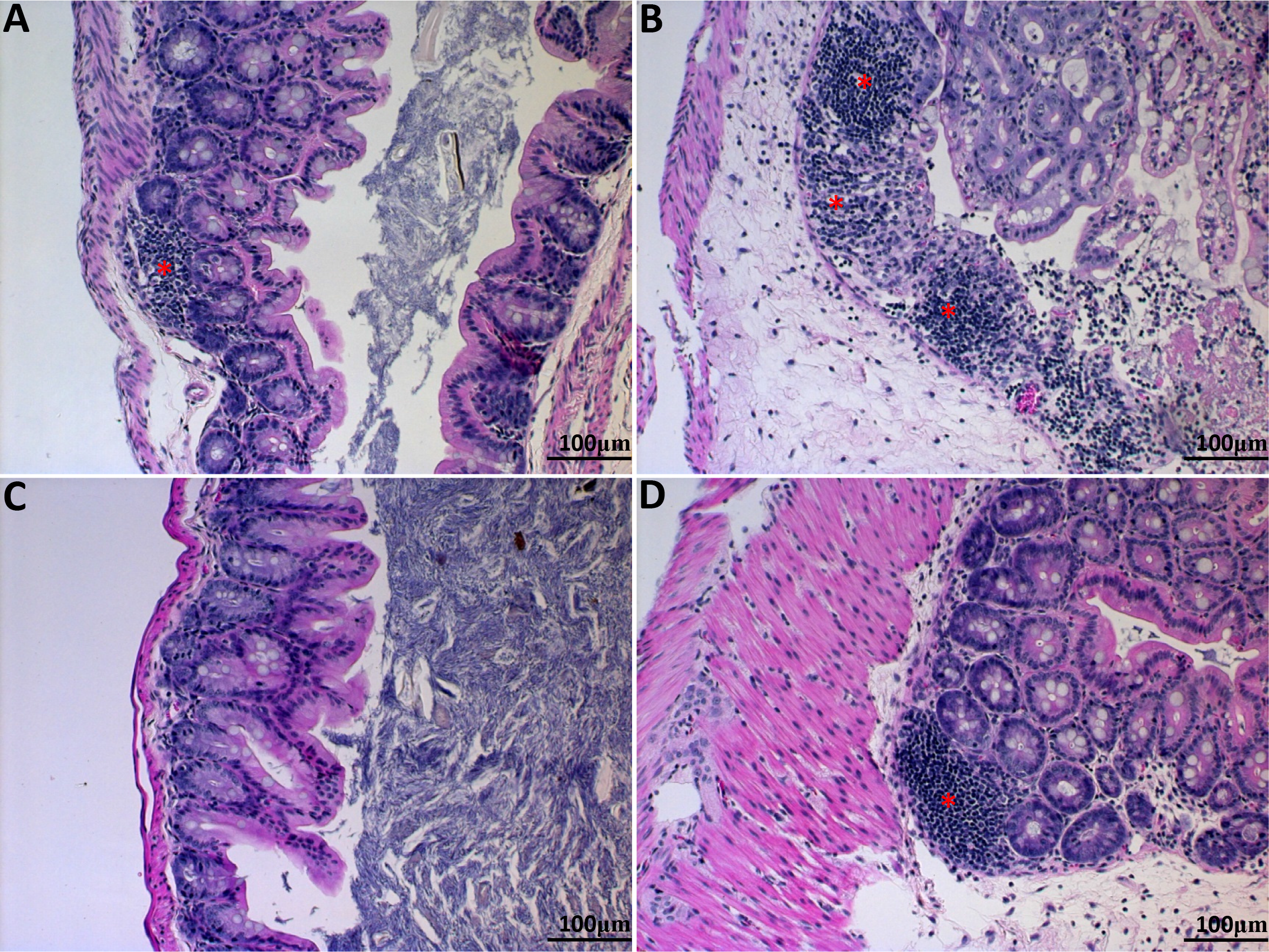
| H&E representative pictures of ILFs in cecum. WT control (A), WT DSS (B), GPR4 KO control (C), and GPR4 KO DSS (D). 20× microscope objective. Red asterisk indicates ILF.

**SI: 3.**
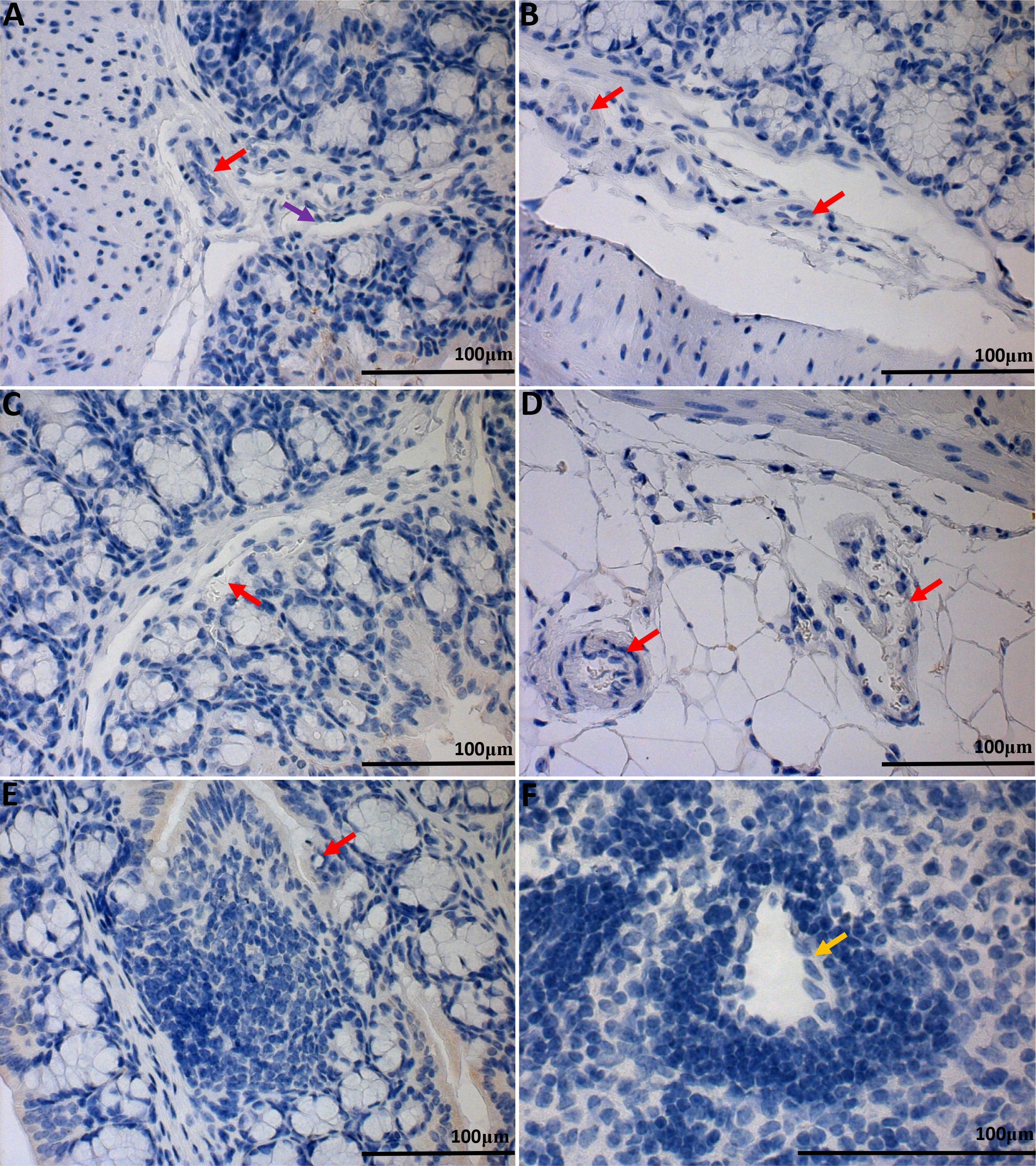
| GFP immunohistochemistry of WT untreated control colon. No visible GFP expression can be detected in tissues. Minor background staining could be observed on epithelium. Submucosa (A&B), transverse folds (C), ex-mural (D), isolated lymphoid follicles (E), and mesenteric lymph node HEV and resident macrophages (F). 40× and 63× microscope objectives. Red arrows indicate blood vessels, yellow arrows indicate HEVs, and purple arrows indicate lymphatics.

**SI: 4.**
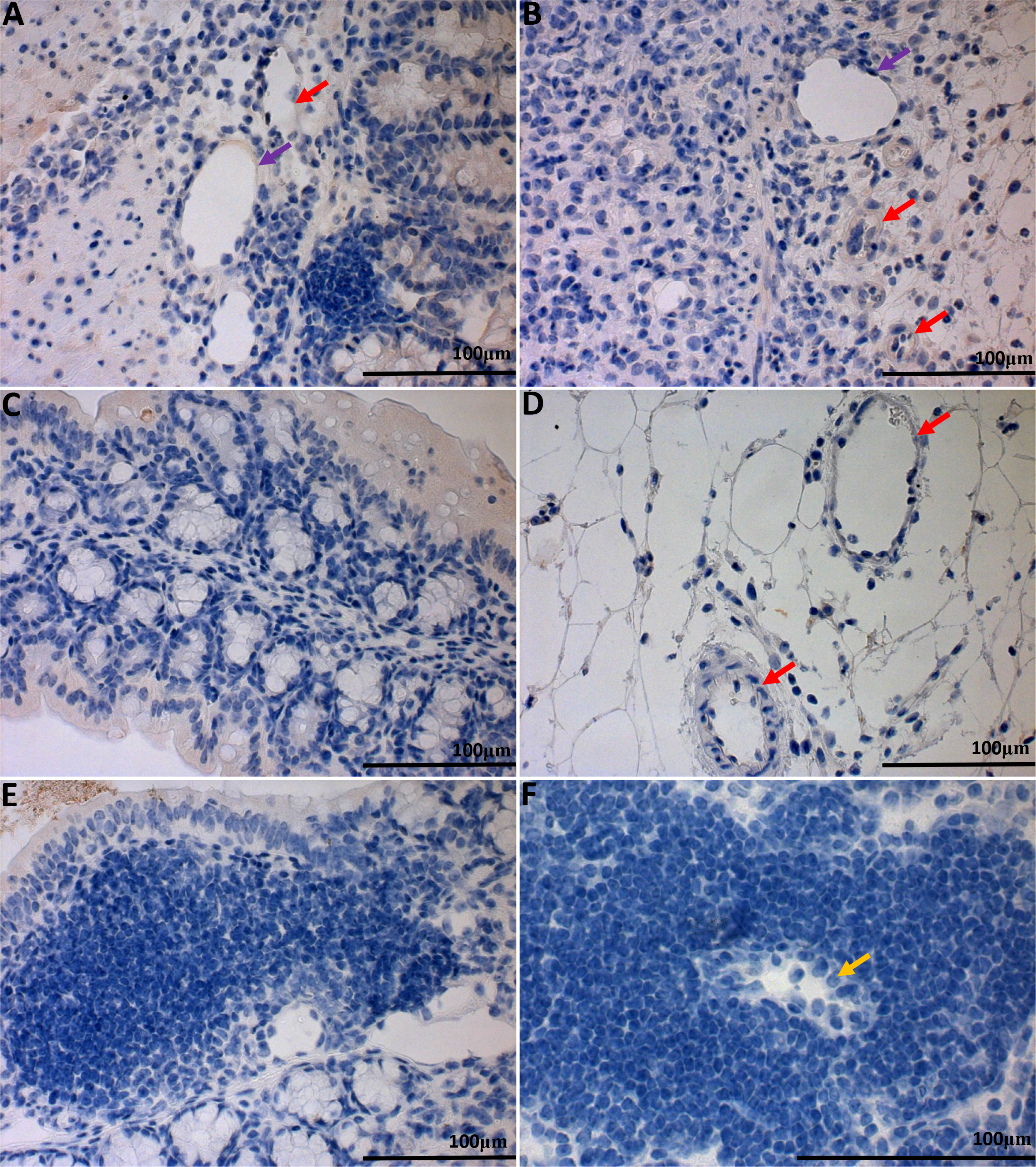
| GFP immunohistochemistry of WT-DSS treated colon. No visible GFP expression can be detected in tissues. Minor background staining could be observed on epithelium, luminal content, and connective tissues. Submucosa (A&B), transverse folds (C), ex-mural (D), isolated lymphoid follicles (E), and mesenteric lymph node HEV and resident macrophages (F). 40× and 63× microscope objectives. Red arrows indicate blood vessels, yellow arrows indicate HEVs, and purple arrows indicate lymphatics.

**SI: 5.**
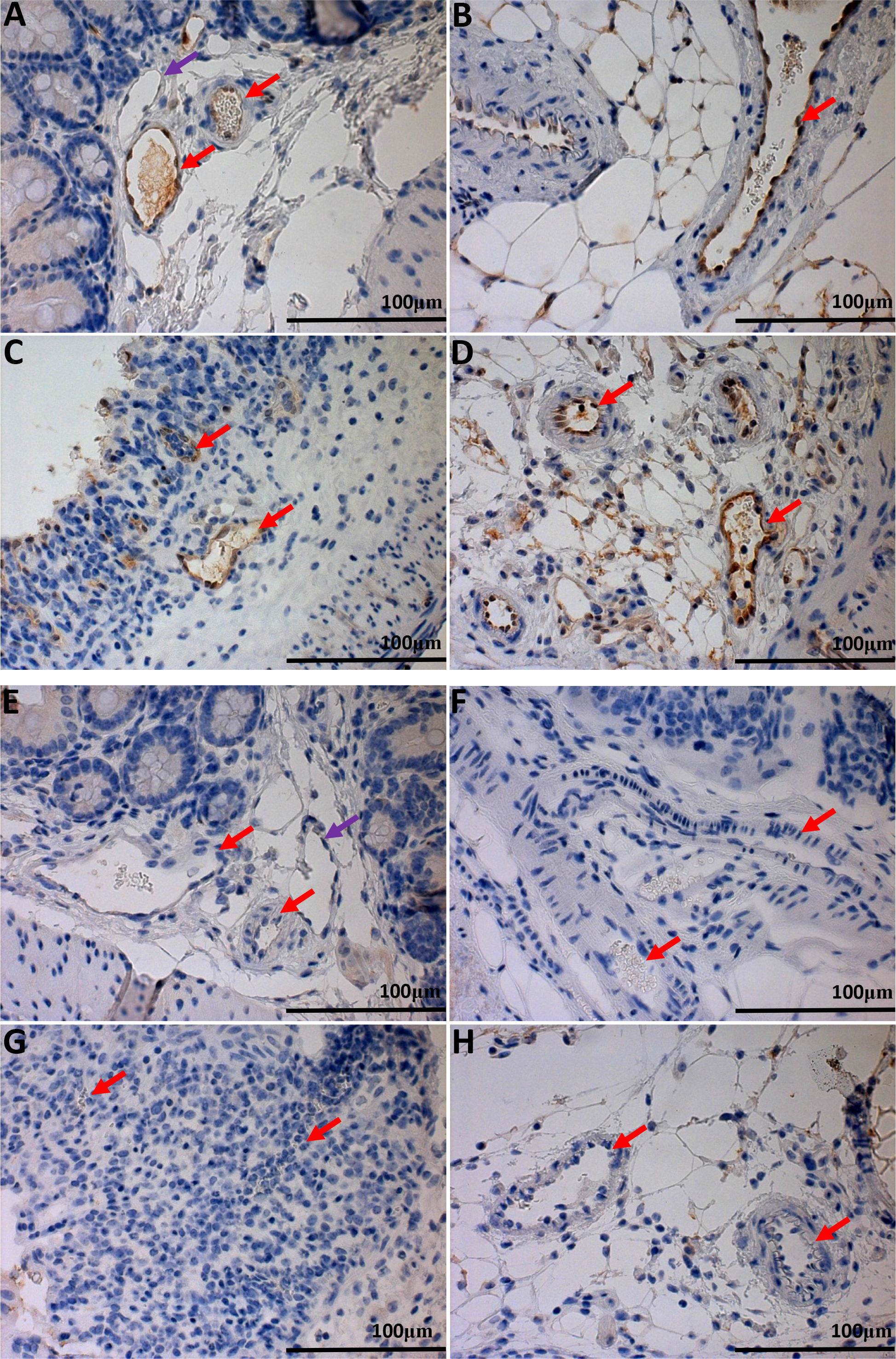
| GFP knock-in as a surrogate marker for GPR4 expression in GPR4 KO mouse cecum. We performed IHC of GFP in GPR4 KO cecum tissue. Similarly as mouse colon tissue, GFP expression could be visualized in the intestinal microvascular endothelial cells, ex-mural blood vessels, arteries in both control and inflamed cecum tissues. Lymphatics were negative for GFP expression. GPR4 KO untreated cecum submucosa blood vessels (A) and ex-mural vessels (B) compared to inflamed GPR4 KO-DSS submucosa blood vessels (C) and ex-mural vessels (D). No GFP signal could be detected in WT control untreated cecum tissues (E&F) and WT-DSS cecum tissues (G&H). 40× microscope objective. Red arrows indicate blood vessels, and purple arrows indicate lymphatics.

**SI: Table 1.** | Taqman Primer-probe set name and ID used for real-time PCR analysis.

| Primer-Probe |  |
| --- | --- |
| E-Selectin | Mm00441278_m1 |
| ICAM-1 | Mm00516023_m1 |
| VCAM-1 | Mm01320970_m1 |
| MAdCAM-1 | Mm00522088_m1 |
| COX-2 (PTGS2) | Mm00478374_m1 |
| CXCL2 | Mm00436450_m1 |
| GPR4 | Mm00558777_s1 |
| 18S rRNA | Hs99999901_s1 |
| β-actin | Hs99999903_m1 |

